# Interplay between Brownian motion and cross-linking kinetics controls bundling dynamics in actin networks

**DOI:** 10.1101/2021.09.17.460819

**Authors:** Ondrej Maxian, Aleksandar Donev, Alex Mogilner

**Affiliations:** Courant Institute, NYU, New York, NY 10012; Department of Biology, NYU, New York, NY 10012

## Abstract

Morphology changes in cross-linked actin networks are important in cell motility, division, and cargo transport. Here we study the transition from a weakly cross-linked network of actin filaments to a heavily cross-linked network of actin bundles through microscopic Brownian dynamics simulations. We show that this transition occurs in two stages: first, a composite bundle network of small and highly aligned bundles evolves from cross linking of individual filaments; second, small bundles coalesce into the clustered bundle state. We demonstrate that Brownian motion speeds up the first stage of this process at a faster rate than the second. We quantify the time to reach the composite bundle state and show that it is a strong function of mesh size only when the concentration of cross links is small, and that it remains roughly constant if we decrease the relative ratio of cross linkers as we increase the actin concentration. Finally, we examine the dependence of the bundling timescale on filament length, finding that shorter filaments bundle faster because they diffuse faster.

## 1 Introduction

The structure and mechanical properties of eukaryotic cells are largely controlled by the actin cytoskeleton, which contains a network of actin filaments interconnected by protein cross linkers (CLs) [1, 6]. Changes in cell mechanical properties, from more viscous to more elastic, relate to corresponding cytoskeletal morphology changes, from a weakly cross-linked network of actin filaments to a network of clustered bundles [34, 28]. The formation of a clustered bundle state has previously been observed in actin suspensions with CLs such as filamin [40, 19], scruin [14], and *α*-actinin [29, 45]. In all of these systems, increasing the concentration of the cross-linking protein progressively transitions the steady state from a homogeneous meshwork, where filaments are distributed isotropically, through a composite bundle state, where bundles are composed of only a few filaments, to the clustered bundle state, where bundles can be separated by distances as large as 100 *μ*m [40, 29].

Usually, the bundled network steady state is the result of a balance between cross-linking and other mechanisms that break up bundles. Indeed, in our previous work [34], we introduced actin filament turnover (to model (de)polymerization) and found that the steady state network morphology is the result of a competition between actin bundling and actin turnover. In particular, we observed either a homogeneous filament meshwork or network of bundles embedded in the filament meshwork, depending on the relationship between the turnover time and the timescale of filament bundling. In most of this paper, we will disable filament turnover and study how the timescale of bundling, which we define approximately as the time to reach the composite bundle state, is affected by the underlying microscopic parameters and the Brownian motion of the filaments. Quantifying this timescale is important because its competition with filament turnover rate determines the steady state network structure, as we will demonstrate in Section 3.5.

While it was observed over 30 years ago [19] that Brownian motion drives bundle formation, its precise mechanism for doing so remains unclear. For instance Hou et al. [19], speculated that rotational diffusion aids in bundling, as filaments that are linked at one location rotate until other locations can be linked together, resulting in a bundle. More recently, it was shown that bundling is most efficient in a fluid-like environment, where actin filaments can diffuse more readily [11, 12]. At minimum, these studies imply that bundling is more difficult without Brownian motion, but could actin filaments still arrange into bundles without it?

The importance of Brownian motion in bundling can be seen in experiments where filament length varies, or when polymerization and bundling are initiated simultaneously. In this case, shorter filaments form a more stable clustered bundle state [22, 12], with the shortest filaments organizing into spindle-type structures [31, 46]. In systems where polymerization and bundling happen simultaneously, it has been shown that the formation of the clustered bundle state can be prevented via an increase in the actin polymerization rate [11]. Mean field theory and simulations show that the slow down in bundling at high polymerization rates could be driven by a combination of steric interactions and the Brownian motion of the fibers being constrained by cross linkers [12]. It remains unclear, however, to what extent the attenuation of bundling is driven by sterics vs. cross linking, and even if a composite bundle state can form if the length of the filaments is larger than the initial mesh size.

An underlying assumption in conceptual explanations of bundling is that sufficient CL is available to cross link filaments once they move closer together. The literature is conflicted, however, on exactly how much CL is sufficient. For instance, in the same experimental system of filamin and actin, some authors report a constant ratio of CL to actin necessary for bundling [19, 45], while others report that the relative amount of CL necessary for bundling decreases as actin concentration increases [40]. There is also a nontrivial effect of temperature on the amount of CL required for bundling; with higher CL-to-actin ratios bundling can occur at lower temperatures [43]. Experimental investigation of the precise amount of CL necessary for the clustered bundle state to form is difficult since the observation of bundles is a qualitative phenomenon with a subjective definition, and therefore varies based on the tools used. Simulations can provide a more definitive analysis of how bundling depends on CL concentration.

Two simulation approaches have been used to theorize about the bundling of actin filaments. One of them was to use equilibrium thermodynamics to find conditions at which the free energy, consisting of translational and rotational entropy of rod-like filaments and enthalpy and entropy of the CL distribution, is lower in the bundled state than in the unbundled mesh [4, 52]. The important results of these theories were that a critical CL concentration is needed for the bundling phase transition and that ultimately one giant bundle has to form, but transiently the filaments could be kinetically trapped in multiple bundles [4, 24]. However, actin bundling is not taking place in thermodynamic equilibrium, and several modeling studies harnessed the Brownian dynamics approach. One of the earliest [50] of these studied the roles of translational and rotational diffusion in bundling of uniformly laterally attracting filaments. A very detailed model in three dimensions in the presence of polymerization, steric interactions and angular stiffness of the filament-CL bond [25] revealed how the morphology of the bundled network scales with mechanical and biochemical parameters. Last, but not least, a combination of scaling estimates and Brownian dynamics simulations with simplified CL properties revealed multi-scale transitions from the isotropic to bundled phase [12]. Most of these previous studies focused on the actin network structure, rather than on the temporal evolution of the bundled state.

In this paper, we use agent-based simulations to quantify the evolution of the clustered bundle state from a homogeneous meshwork of filaments and examine the role of Brownian motion therein. We begin in Section 2 by describing our computational methods [33, 34]. In Section 3, we demonstrate how a composite bundle state, and subsequently a clustered bundle state, evolve from a homogeneous meshwork, similar to what is observed in experimental networks [28]. We introduce a timescale, *τ_c_*, that quantifies the time to reach the composite bundle state, and show that the dynamics on shorter, but not so much on longer, timescales are accelerated by Brownian motion. While we do not consider steric interactions, we demonstrate that the strong cross linking present at later times is sufficient to arrest the bundling process. We also show that the bundling timescale is limited by filament diffusion for smaller CL concentrations, while for larger CL concentrations this diffusion has a minor effect. We find that the relative CL-to-actin ratio required to achieve the same bundling time decreases with increasing actin network mesh size. Finally, we show that the diffusion effect explains the faster bundling for shorter filaments. We discuss some remaining questions, and possible extensions of our model necessary to answer them, in Section 4.

## 2 Methods

We begin with a review of the kinematics of inextensible fibers, slender body hydrodynamics, and our model of dynamic cross linking [33, 34] in Section 2.1. We then discuss new material pertinent to the simulation of actin bundles, including how we modify our algorithm to simulate rigid fibers (Section 2.2) and to account for their translational and rotational diffusion (Section 2.3). Once we introduce thermal motion, a consistent model also requires us to keep the CL dynamics in detailed balance, i.e., that the binding and unbinding dynamics are in equilibrium with respect to the Gibbs-Boltzmann distribution. We account for this in Section 2.4 via a simple change to the rates of CL binding. Finally, in Section 2.5, we discuss how we use a time-splitting algorithm to evolve the system in time. While the CL binding and unbinding dynamics and filament evolution are treated in a first-order-accurate manner, we use a higher-order integrator for the Brownian term that can more accurately preserve fluctuation-dissipation balance.

### 2.1 Dynamic cross linking of semiflexible, inextensible fibers

This section reviews our algorithm for simulating the dynamic cross linking of semiflexible fibers [33, 34], beginning with the kinematics of inextensible fibers and slender body hydrodynamics [33], and concluding with our model of dynamic cross linking [34]. As in our previous work [33, 34], we use a periodic boundary condition in all three dimensions to mimic a bulk suspension.

#### 2.1.1 Semiflexible, inextensible fibers

We represent the centerline of each fiber by the Chebyshev interpolant ***X***(*s*), where *s* ∈ [0, *L*] is arclength and *L* is the fiber length. Likewise, the corresponding fiber tangent vector is represented by *τ*(*s*) = ***X***_*s*_(*s*). Because the fibers are inextensible, the tangent vector should have unit length for all time, ***τ***(*s*, *t*) · ***τ***(*s*, *t*) = 1, for all *s* and *t*. Differentiating this constraint with respect to time, we obtain ***τ***_*t*_ · ***τ*** = 0, so that the velocity of the filament centerline can be parameterized as [33]

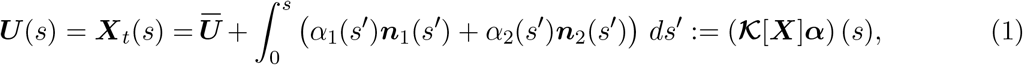

where ***τ***(*s*), *n*_1_(*s*), *n*_2_(*s*) are an orthonormal coordinate system at each *s*, and *α*_1_(*s*) and *α*_2_(*s*) are two unknown functions. Equation (1) defines a continuum kinematic operator 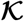 that parameterizes the space of inextensible motions [33, Sec. 3].

To close the system and solve for 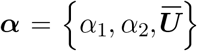, we need to state the forces acting on the fiber centerline. To enforce the inextensibility constraint, we introduce a Lagrange multiplier force density ***λ***(*s*, *t*). In addition to the constraint force, the fibers are also subject to a bending force with density ***f^κ^*** [***X***] = -*κ**X**_ssss_*, where *κ* is the bending stiffness, and an external force density that comes from any attached cross links, which we denote by ***f***^(CL)^. The total force density at every instant in time is therefore ***f*** = ***λ*** + ***f*^*κ*^** + ***f***^(CL)^. Introducing the hydrodynamic mobility operator 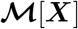 that gives velocity from force (density), the evolution equation of the fiber centerline can be written as

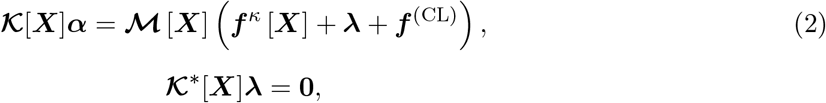

subject to the “free fiber” boundary conditions [44]

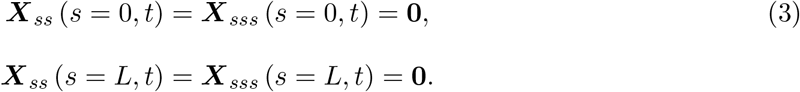

We solve (2) for the kinematic coefficients ***α*** and constraint forces ***λ***. The adjoint condition 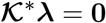 closes the system of equations, and encodes the principle of virtual work that constraint forces ***λ*** do no work for any inextensible motion of the fiber centerline [33, Sec. 3.4]. We still have to discuss the evaluation of 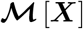 and ***f***^(CL)^, which we do next.

#### 2.1.2 Mobility evaluation

In previous work [33, 34], we utilized three different approaches to evaluate the mobility operator 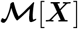. All of these approaches are based on traditional slender body theories [23, 21], which relate the velocity of a slender filament in Stokes flow to the force density exerted on its centerline. The total velocity at a point on the filament can be broken into three parts: that from force concentrated near the point (the “local drag” part, which dominates as the fiber becomes infinitely slender), that from the rest of the filament (intra-fiber hydrodynamics), and that from forcing on other filaments (through hydrodynamic interactions mediated by the fluid medium). The first two of these are simple to evaluate, given that they can be computed on each filament separately, but the third is expensive to compute because it involves all-to-all interactions through the fluid.

We have already studied the role of nonlocal hydrodynamic interactions in previous work [34], where we found that the time required to reach a particular bundled state is underestimated by at most 10–20% when inter-fiber hydrodynamic interactions are dropped. In this paper, our interest will be in how parameters other than hydrodynamic interactions affect the bundling time. Therefore, to improve computational efficiency, we will ignore hydrodynamic interactions between distinct filaments and evaluate the mobility by including only the local drag part and intra-fiber hydrodynamics. Specifically, the mobility operator on each fiber is given by nonlocal slender body theory [23, 21]

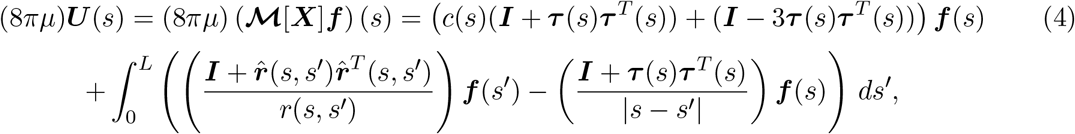

where ***r***(*s*, *s*′) = ***X***(*s*) – ***X***(*s*′), *r* = ||***r***||, 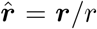, and *c*(*s*) is a local drag coefficient which has a logarithmic dependence on the fiber radius *a*. Away from the fiber endpoints, we use the classical result [23]

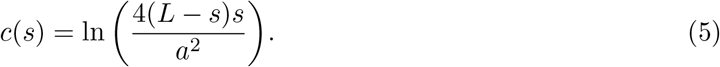

Near the endpoints, we regularize (5) over a distance *δ* = 0.1*L* as discussed in [33, Section 2.1]. The choice of mobility (4) allows us to simulate the evolution of bundles faster, and prevents possible numerical problems that could result when evaluating the nonlocal flows induced by hundreds of filaments in a bundle on each other [34].

#### 2.1.3 Evaluation of *f*^(CL)^

We use a stochastic simulation algorithm to update the locations of the dynamic cross linkers. At each time step, this algorithm, which we discuss in the next section, gives the fiber indices *i* and *j* that are linked by each link, as well as the arclength coordinates 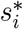 on fiber *i* and 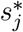 on fiber *j* where the link is bound. Letting *K_c_* be the link stiffness (units force/length) and *ℓ* the CL rest length, we define the force density on fiber *i* due to the CL as

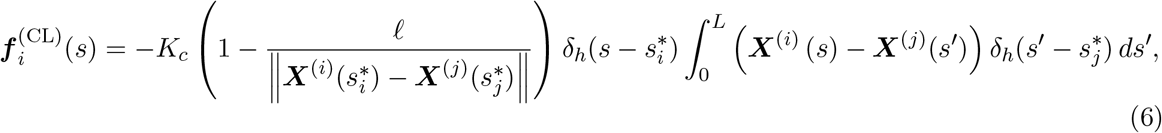

where *δ_h_* is a Gaussian density with standard deviation *σ*. While *σ* → 0 corresponds to a standard spring point force, we use a finite *σ* to preserve smoothness for our spectral numerical method. For *N* = 16 points per fiber, which we use throughout this paper, we use *σ*/*L* = 0.1 [33]. As discussed in [34], this model is an approximation to the complex elasticity of *α*-actinin, and is based on experimental observations that the torsional stiffness of the *α*-actinin-actin bond does not influence the dynamics of that bond [7].

#### 2.1.4 Dynamic cross-linking

Our model of dynamic cross linking is discussed in detail in [34]. Briefly, we discretize each fiber into *N_u_* uniformly-spaced “binding sites” with distance Δ*s_u_* = *L*/(*N_u_* – 1) between the sites. We make the assumption that the diffusion of individual CLs is sufficiently fast that it can be coarse-grained into a single binding rate *k*_on_ with units 1/(length×time). This means that a CL end can bind to a single discrete fiber binding site with rate *k*_on_Δ*s_u_* per second. In the absence of Brownian motion, as in [34], when one end of the CL is bound, the second end can bind to a nearby fiber with rate *k*_on,s_. By “nearby” we mean a binding site on a distinct fiber that is within a distance interval

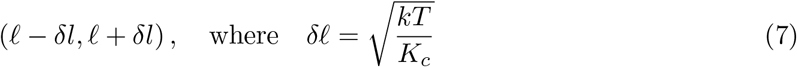

from the first bound end, where *ℓ* is the CL rest length and *δℓ* is a measure of the fluctuations in spring length.

Each of the binding reactions has a reverse reaction: a CL with both ends bound can have one end unbind, leaving one end bound, with rate *k*_off,s_, and a CL with one end bound can unbind with rate *k*_off_ to have zero ends bound. There are thus four possible reactions, which we simulate stochastically using a version of the standard Stochastic simulation / Gillespie algorithm [15, 2]. The details of our implementation can be found in [34].

In the clustered bundle states that we simulate here, the number of links attached to a given site can grow without bound. To prevent this, we introduce a CL width *c_w_* = 20 nm [36] and set the maximum number of bound CLs at each site to ⌈Δ*s_u_*/*c_w_*⌉. We implement this in the stochastic simulation algorithm using rejection: if a binding event is selected and the binding site is full, we simply move on to the next possible event.

### 2.2 Modifications for rigid fibers

In order to straightforwardly account for thermal fluctuations, we will consider the case when the fibers are rigid, so that the only possible fluctuations are translational and rotational diffusion. To simulate rigid fibers, we modify the kinematic operators 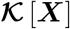 and 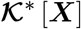 in (1) and (2). For rigid fibers, we introduce ***α*** ≡ ***V*** = {***U**_c_*, **Ω**}to parameterize the space of rigid body motions, where ***U**_c_* = *d**X**_c_*/*dt* is the translational velocity of the fiber center ***X***_*c*_ = ***X***(*L*/2), and **Ω** is the angular velocity of the fiber about its center. This gives the fiber velocity

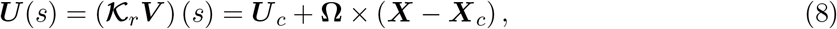

which reduces the constraint of virtual work to the fact that ***λ*** produces no net force and torque,

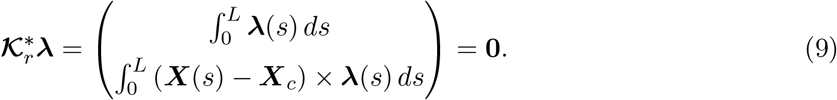

We then solve the system (2) with 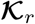 and 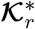 replacing 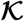 and 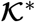. In the Supplementary Text, we show how to easily generalize our discretization for inextensible fibers [33] to straight rigid fibers by restricting the number of Chebyshev modes included in the kinematic operator 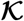 to only the first one.

Because the fibers are rigid, we can formulate the hydrodynamic mobility as a 6 × 6 mobility matrix ***N***[***X***] which computes the fiber motion due to a total force ***F*** and torque ***T***,

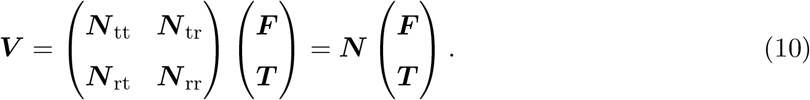

When the fibers are straight, as in this work, and we measure the mobility about the geometric center of the fiber, the cross translation-rotation and rotation-translation mobilities vanish, ***N***_tr_ = ***N***_rt_ = **0**. We recall that in this work we neglect hydrodynamic interactions between fibers, so the mobility matrix ***N*** can be computed for each fiber separately.

The mobility ***N*** can be obtained numerically from the slender body mobility matrix ***M*** (see [33, Sec. 4.2] for the discretization) via the Schur complement [42, 47]

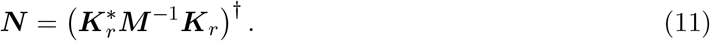

Note that the pseudo-inverse is required because applying a torque about the axis of a straight fiber produces no net motion (other than twisting, which we do not account for here). For straight fibers with constant tangent vector ***τ***, by symmetry, the mobility ***N*** must be of the form

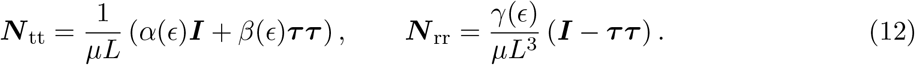

In Supplementary Table S1, we tabulate the coefficients *α*, *β*, and *γ* for biologically-relevant *ε*. See also [51] for semi-analytical approximations.

### 2.3 Thermal fluctuations with rigid fibers

For Brownian dynamics simulations, we need to solve the overdamped Ito Langevin equation

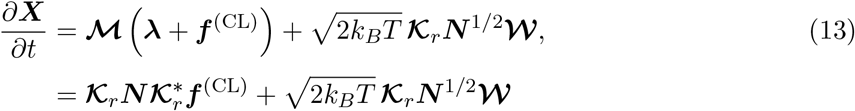

where 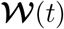 is a vector of six i.i.d white-noise processes, and ***N***^1/2^(***N***^1/2^)^*T*^ = ***N***. The last equality, which puts the overdamped Langevin equation into the more traditional symmetric form, follows from the fact that the deterministic velocity can be written using (9) and (10) as

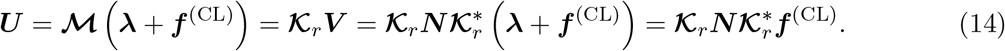

Note that because the fiber mobility is measured around the obvious geometric center, there is no stochastic drift term in the resulting Ito overdamped Langevin equation (13) [32, 8]. In our Brownian dynamics simulations with straight fibers, we use the precomputed values of *α*, *β*, and *γ* in (12) to generate 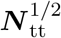 and 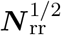 from the fiber tangent vector ***τ*** according to (S10).

The random displacement of a fiber over a time interval *τ* can be sampled the following way,

1. Draw a vector ***W*** of six i.i.d. standard Gaussian variates and sample the rigid velocity

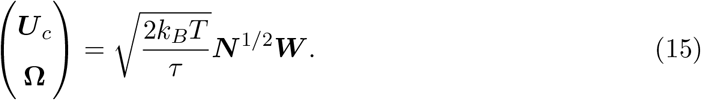
2. Update the fiber by translating its center by ***U****_c_τ* and rotating the fiber about its center by an oriented angle **Ω***τ*.

Note that for straight fibers one can simplify the formulation of Brownian dynamics; the formulation presented here applies to curved rigid fibers as well.

### 2.4 Keeping the CL dynamics in detailed balance

When we account for thermal translation and rotation of the fibers, we also want to be sure that the binding and unbinding of the links is consistent with detailed balance, which is not the case for the constant rates we introduced in Section 2.1.4. Let ***C*** denote a configuration of *C* links (list of fiber pair connections), and ***x*** denote the configuration of fibers (binding site positions). The desired Gibbs-Boltzmann equilibrium distribution is

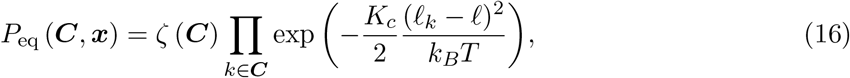

where *ℓ_k_* is the length of link *k*, and *ζ*(*C*) determines the probability to observe the cross-linking configuration ***C***. Now consider a transition to/from a state ***C*′** with one added link ***C*′**, which has length 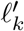. Then at equilibrium the transition between the two states must obey

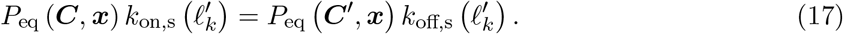

Substituting *P*_eq_ from (16) into (17), we obtain the constraint of detailed balance

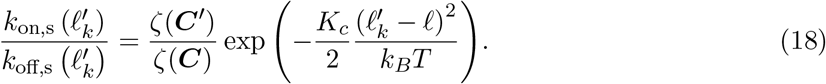

To satisfy (19) for every choice of ***C*** and ***C***′ with binding and unbinding rates that only depend on spring length and not ***C*** and ***C*′**, we must have

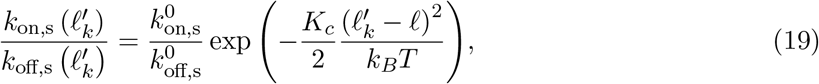

where 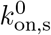 and 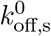 are the transition rates for a link at rest length.

To satisfy (19), we maintain a constant 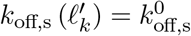 and set

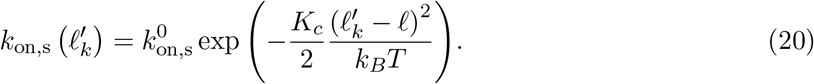

Other choices are possible; for example, the rate of unbinding can depend on the stretch [3, 10]. To efficiently search for possible binding pairs, we approximate the set of all binding combinations by setting the maximum link stretch in (7) to be two standard deviations of the Gaussian (20), i.e., 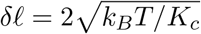.

### 2.5 Temporal integration

We employ a time splitting approach to evolve the cross-linked actin network. At each time step, we have three processes to simulate: the thermal diffusion of the fibers, the binding and unbinding of the dynamic CLs, and the deterministic evolution of the fiber positions. The last two of these steps are laid out in full in [34], where we employ Lie splitting to first process binding and unbinding events, and then use the method developed in [33] to evolve the fiber positions in an inextensible (or rigid, see Supplementary Section 5.1) way. Here we use the first-order accurate, backward Euler version of the deterministic fiber update which is discussed in [34].

It remains to determine how we will treat the Brownian update. We use a second-order Strang-type splitting scheme, where during each time step of duration Δ*t* we:

1. Randomly displace and rotate the fibers over a time interval *τ* = Δ*t*/2 using the algorithm in Section 2.3.
2. Update the cross link attachments (using the stochastic simulation algorithm) and perform a deterministic fiber update, both over a time interval Δ*t*, using the method of [34].
3. Randomly displace and rotate the fibers over a time interval *τ* = Δ*t*/2 using the algorithm in Section 2.3.

### 2.6 Network statistics

We quantify the evolution of the cross-linked actin network by examining the connectivity of the fibers in two ways. First, given the total number of CLs in the system *C*, we compute an average link density per fiber via the formula “Link density”= 2*C*/(*LF*). Second, we map the network to a connected graph to study how the structure evolves in time [31]. We define a “bundle” as a connected group of at least *FB* = 2 filaments, where a connection between two fibers is a pair of links with anchoring locations at least *d*_bund_ = *L*/4 apart on each fiber [34], so that the links limit the fibers’ rotational degrees of freedom. We then define two measures of the degree of bundling in the system. The first measure is the bundle density, which is the number of bundles per unit volume 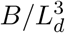, where *B* is the number of bundles and *L*_d_ is the length of the simulation cell. The second measure is the percentage of fibers in bundles, defined as the percentage of filaments connected to at least one other filament by two links at least *d*_bund_ = *L*/4 apart. The bundle density statistic preferentially weights smaller bundles, since a bundle of two filaments is counted the same as a bundle of five filaments, while the percentage of fibers in bundles is independent of *F_B_*.^1^

For a bundle of *b* filaments, we define an orientation parameter as the maximum eigenvalue of the matrix [31]

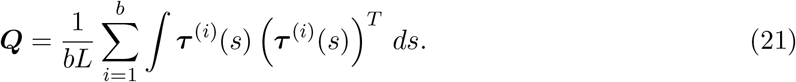

The orientation parameter takes values in [1/3,1], with 1 being the value for a group of straight fibers with the same tangent vector. Given information about the bundles, we compute an average bundle orientation parameter by taking an average over bundles with at least two filaments, weighted by the number of filaments in each bundle.

Throughout this paper, we will quantify the concentration of fibers in terms of the initial mesh size [37] of the suspension, 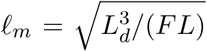 (parameters are defined in Table 1). Note that this estimate for *ℓ_m_* applies to non-bundled (disordered) suspensions of fibers, so really when we use *ℓ_m_* we mean the *initial* mesh size, prior to the bundling process beginning. We will operate in the regime where the fluctuations in the CL rest length as defined in (7), which are of magnitude *δℓ* = 20 nm (see parameters in Table 1), are several times smaller than the typical filament spacing, which is at most the initial mesh size 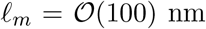 and at least the cross linker length of 50 nm.

**Table 1:** Simulation parameters.

| Parameter | Definition | Value | Unit | Notes |
| --- | --- | --- | --- | --- |
| $a$ | Fiber radius | 4 | nm | [18] |
| $L$ | Fiber length | 0.5, 1 | $\mu\text{m}$ | [20, 35] |
| $F$ | Number of fibers | 200 – 1600 | | |
| $L_d$ | Simulated volume’s extent | 2 – 4 | $\mu\text{m}$ | Cubic unit cell |
| $\mu$ | Cytoplasm viscosity | 0.1 | $\text{Pa}\cdot\text{s}$ | 100× water (cytoplasm) [30] |
| $\kappa$ | Fiber bending stiffness | 0.07 | $\text{pN}\cdot\mu\text{m}^2$ | 17 $\mu\text{m}$ persistence length [16] |
| $K_c$ | CL spring stiffness | 10 | $\text{pN}/\mu\text{m}$ | [27] |
| $\ell$ | CL rest length | 50 | nm | [36] |
| $k_{\text{on}}$ | CL first end binding rate | 5 | $1/(\mu\text{m}\cdot\text{s})$ | [34] |
| $k_{\text{on},s}$ | CL second end binding rate | 50 | $1/(\mu\text{m}\cdot\text{s})$ | $k_{\text{on},s} \gg k_{\text{on}}$, not measured |
| $k_{\text{off}}$ | CL (one end bound) unbinding rate | 1 | 1/s | [26, 48] |
| $k_{\text{off},s}$ | CL (both ends bound) unbinding rate | 1 | 1/s | $k_{\text{off},s} = k_{\text{off}}$ |
| $c_w$ | Actin binding site width | 20 | nm | [36] |
| $k_B T$ | Thermal energy | $4 \times 10^{-3}$ | $\text{pN}\cdot\mu\text{m}$ | 25°C |
| $N$ | Number of Chebyshev points | 16 | | [33, Sec.6.3.1] |
| $\Delta s_u$ | Binding site spacing | 0.026 | $\mu\text{m}$ | [34] |
| $\Delta t$ | Time step size | $10^{-4}$ | s | Limited by $K_c$ |

## 3 Results

We begin this section by discussing the kinetics of bundling for *non-Brownian*, *semiflexible* fibers, and establish that semiflexible fibers with a persistence length similar to that of actin [16] can be well-approximated by rigid filaments. We then show that rigid, Brownian fibers have similar kinetic behavior, except that the Brownian motion (translational and rotational diffusion) speeds up the timescale of bundling.

After these preliminaries, we use our simulations to clarify and explain some of the experimental results on the dynamic formation of cross-linked actin bundles. First, we show how the timescale needed to reach the composite bundle state depends on the fiber concentration (initial mesh size) and concentration of CLs (which controls *k*_on_ in our model). We show that the bundling process is slower when the actin or CL concentration is lower, but that bundling can still occur at low actin concentration, provided there are enough CLs available to bundle the fibers, and that the relative amount of CLs needed for a fixed bundling time decreases as actin concentration increases. Second, we show that the experimental result that bundling occurs faster for shorter fibers [22, 11] can only be reproduced in systems where we consider translational and rotational diffusion. Unless otherwise noted, we will use the simulation parameters listed in Table 1. As discussed in [34], the cross-linking parameters are chosen to mimic *α*-actinin, although we will compare our results to systems with different CLs, such as filamin [40].

### 3.1 Kinetics of bundling for non-Brownian, semiflexible fibers

We begin with simulations in a system of initial mesh size *ℓ_m_* = 0.2 *μ*m, which translates to *F* = 200 filaments in an *L_d_* = 2 *μ*m domain, *F* = 675 filaments in an *L_d_* = 3 *μ*m domain, and *F* = 1600 filaments in an *L_d_* = 4 *μ*m domain. The mesh size we use is of the same order of magnitude as that in cell cortex in vivo [9] and corresponds to 10 to 15 *μ*M G-actin concentration often used in in vitro experiments [29, 45]. In Supplementary Fig. S2, we show that the statistics of the bundling process are insensitive to the domain size up to the point where there is mass coalescence of almost all the fibers in the simulation cell. For this reason, we will consider results from only one set of simulations, the one with *F* = 675 filaments and *L_d_* = 3.

We initialize the set of *F* filaments with random locations and orientations, then during each time step we evolve the fibers by updating the dynamic CLs and then updating the fiber positions in sequential order. Figure 1 shows how the bundling process evolves in small and large systems. On the microscopic scale, filaments that are initially not parallel are linked by CLs, which pull them closer together and allow more links to bind. The binding of additional links leads to the alignment of filaments. Note that the key to the bundling process is the flexibility of the CL, in particular rapid thermal fluctuation of the CL length, which is present implicitly in our model from (7). Because the CLs are small, fluctuations in their length occur on a time scale which is much faster than other characteristic time scales, and so we do not model the fluctuations explicitly. The combination of the CLs’ elasticity and length fluctuations is crucial, as the length fluctuations effectively allow the CLs to “find” the neighboring fibers and bind them, whereupon the elasticity of the CL aligns the fibers, making further cross linking faster.

**Figure 1:**
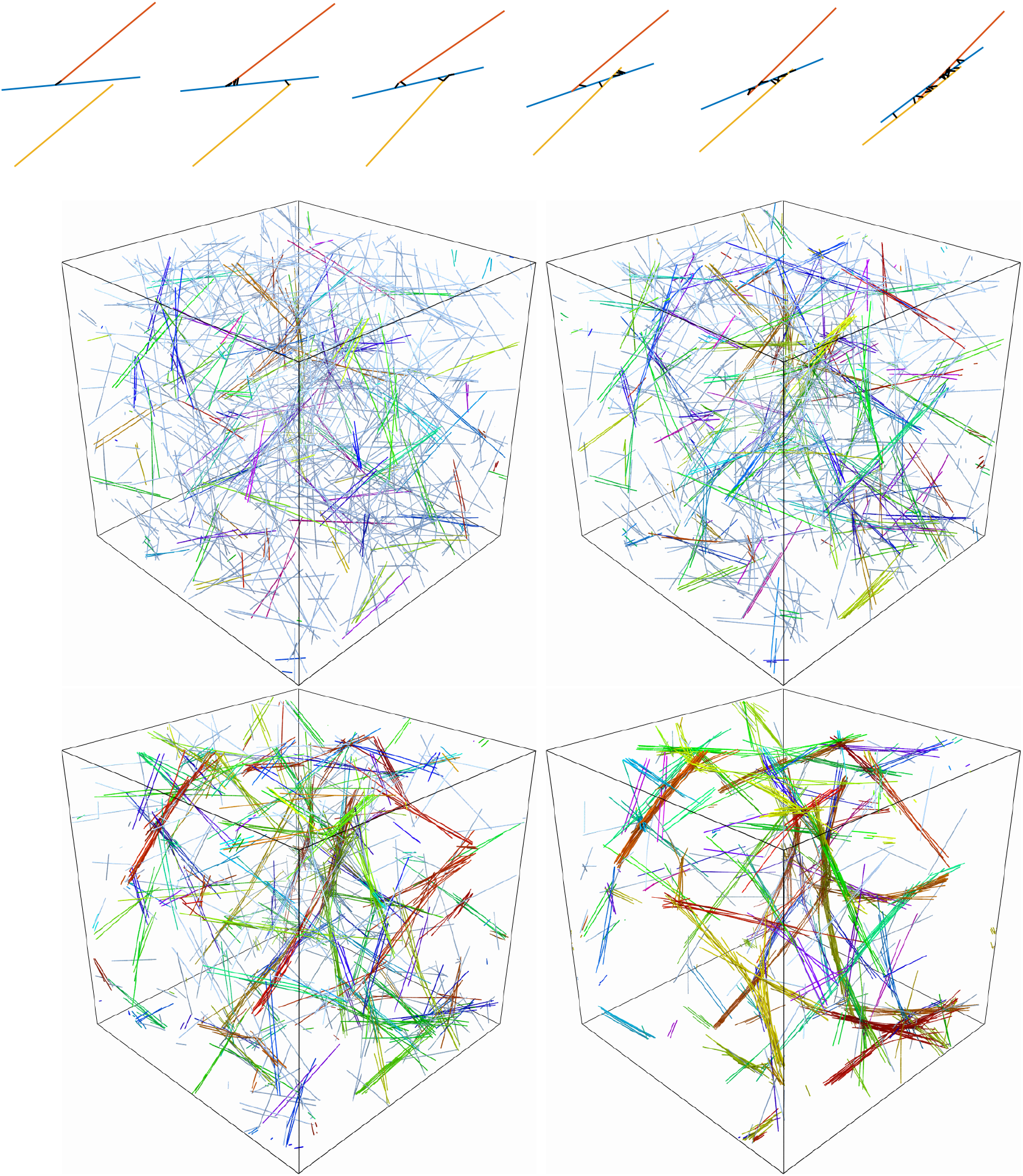
Bundling dynamics on small and large scales. Top: a small-scale bundling process with three filaments and snapshots taken at times *t* = 0, 2, 4, 6, 8, and 10 s. Bottom: Snapshots of the bundling process taken (from left to right and top down) at *t* = 5, 10, 20, and 40 seconds for semiflexible fibers with stiffness *k* = 0.07 pN· *μ*m^2^. Fibers in the same bundle are colored with the same color. The two networks at the middle are before the coalescence transition time *τ_c_* ≈ 16 s, while the two networks at the bottom are after the coalescence time.

This process plays out on a larger scale in snapshots from the simulations, shown in Fig. 1 at *t* = 5,10,20, and 40 seconds. The initial stage of bundling (first two snapshots) is characterized by bundles of a few straight, aligned filaments, which is similar to the experimentally-observed composite bundle state [28] and the three-filament bundle shown at the top of Fig. 1. Later times (bottom two frames) show coalescence of these smaller bundles into larger bundles, with some curvature appearing in the fibers in the final frame. By *t* = 40 seconds, there are only a few bundles made of coalesced smaller bundles, and the network resembles the experimentally-observed clustered bundle state [28], which approaches the low energy state consisting of a single aligned bundle [38].

To quantify our observations, in Fig. 2 we plot the mean link density (2*C*/(*LF*), see Section 2.6), bundle density 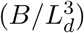, percentage of fibers in bundles, mean bundle alignment parameter, and mean and maximum bundle size throughout the bundling process for three values of fiber stiffness: *κ* = 0.07 (the value for actin), *κ* = 0.007 (fibers ten-fold less stiff), and *κ* = ∞ (rigid filaments). In all systems, we see the number of links per fiber grow in time to approach the maximum of ⌈Δ*s_u_*/*c_w_*⌉ × *L*/Δ*s_u_* = 80, while the bundle density in all systems exhibits a peak around a critical time *τ_c_* ≈ 16 seconds. At this time, the other panels of Fig. 2 tell us that 60% of the fibers are already in bundles, which have a mean alignment parameter larger than 0.9. Figure S3 gives a more precise look at the composition of the bundles, which are the same for the three values of stiffness when *t* ≤ *τ_c_*: at *t* = *τ_c_*, most (> 50%) of the fibers are in bundles of size 11 or less, with a small percentage (< 10%) in bundles of size 10 – 20, and the other 40% of the filaments not in bundles at all. Thus, a time *τ_c_* into the bundling process, most of the fibers are in small, highly aligned bundles, as we see in the snapshots in Fig. 1, and the dynamics up to this point are roughly independent of the fiber stiffness. Based on Fig. 1, we can also think of *τ_c_* as the time required to reach the composite bundle state. For this system of non-Brownian filaments, Fig. 2 shows *τ_c_* ≈ 16 s corresponds to the timescale of increase of the percentage of fibers in bundles (see middle left frame), meaning it is also the timescale on which the fibers’ rotational degrees of freedom are arrested or constrained.

**Figure 2:**
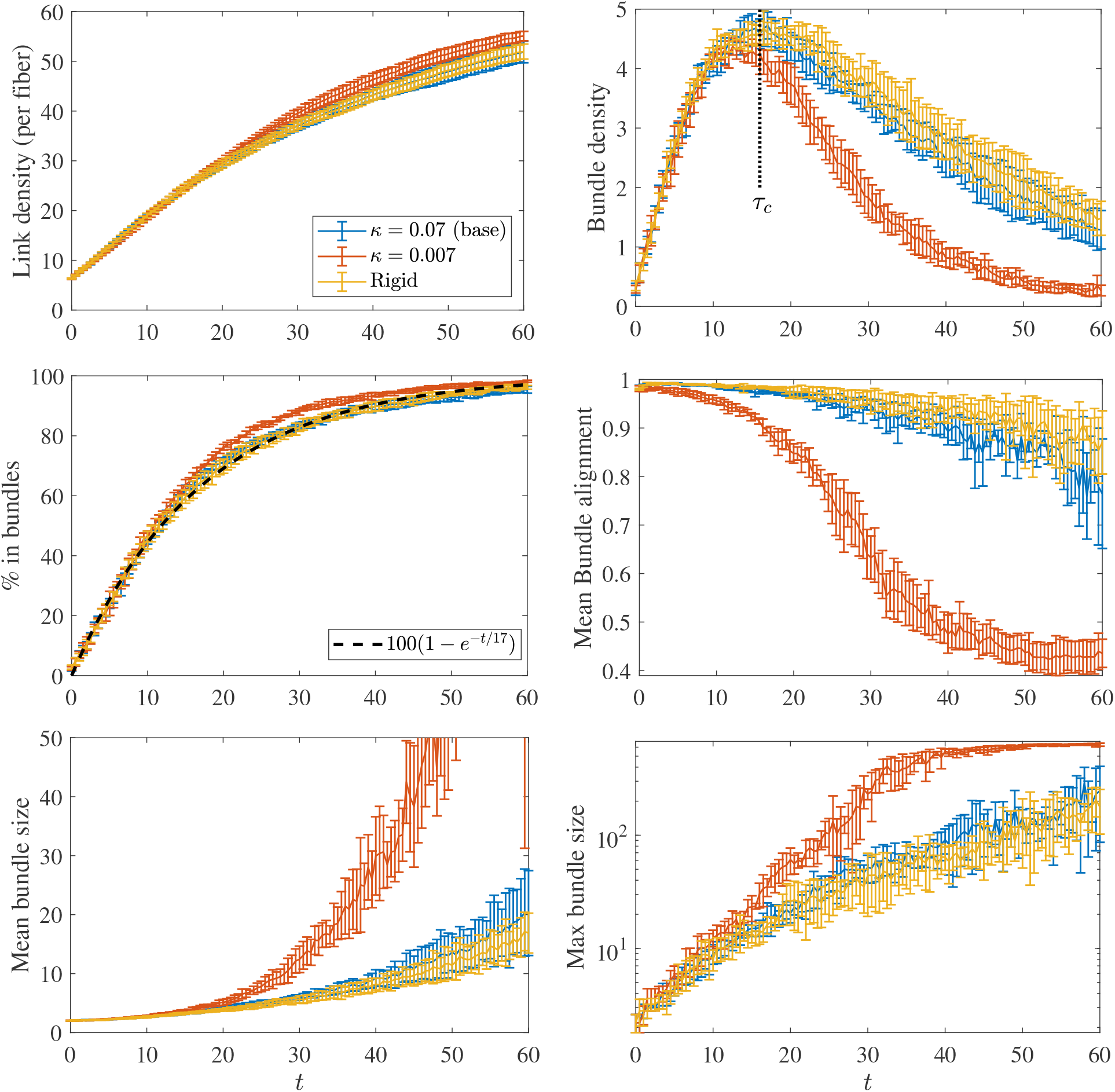
Statistics for the bundling process, where we compare the base parameters (*κ* = 0.07 pN· *μ*m^2^, blue) with the systems with smaller bending stiffness (*κ* = 0.007, orange) and rigid fibers (yellow, *k* → ∞). After *τ_c_* ≈ 16 s, the bundling dynamics for the less stiff fibers are significantly faster. Fibers with similar bending stiffness to actin are well-approximated by rigid fibers.

After the coalesence time, we see a transition to coalescence of bundles, and the flexibility of the fibers comes into play. Figure 2 shows that the number of bundles is declining and the mean bundle alignment is dropping for *t* > *τ_c_*, which implies that bundles are forming with non-aligned fibers. The mean and maximum bundle sizes also start to grow, which again means that small bundles are coming together to form the larger ones we see in the bottom row of Fig. 1. Figure S3 shows that by *t* = 60 seconds, at least 75% of the fibers are in bundles of size 30 or larger. It is in this stage where the flexibility of the fibers can become important: when *κ* = 0.007 (fibers ten-fold less stiff than actin), coalescence of bundles occurs faster than in systems with *κ* = 0.07 or systems with rigid fibers, since in the former case the fibers are more compliant and can be linked together more easily by deforming. That said, when *κ* = 0.07 (persistence length 17 *μ*m), Figs. 2 and S3 show that the dynamics throughout the bundling process are well-approximated by rigid filaments. This analysis is of course limited by the fiber length we’ve chosen: in particular, we have shown that, in the absence of Brownian motion, rigid filaments are a good approximation to semiflexible actin filaments for fibers of length ≤ 1 *μ*m, which are most common in vivo. The approximation will be worse as the filament length gets larger. Henceforth, we will consider rigid fibers only.

### 3.2 Thermal fluctuations speed up the bundling process

We now consider simulations with rigid fibers, for which we can account for translational and rotational diffusion using standard Brownian dynamics methods [32], while maintaining detailed balance in the cross-linking kinetics, as discussed in Section 2.4. An important quantity in this case is the time for a fiber to diffuse across a mesh size. In our initial set-up, the fibers are spaced approximately *ℓ_m_* apart, and they first must find each other in order to cross link and begin the bundling process. The theoretical translational diffusion coefficient of a straight fiber, derived in [32], can be written in terms of the 3 × 3 translational mobility matrix ***N***_tt_ for rigid body motions as

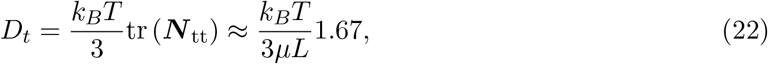

where the last equality gives the result for a fiber aspect ratio of *ε* = 0.004, which we obtain from slender body theory with intra-fiber hydrodynamics (see Table S1, and note that this estimate accounts for the anisotropy of the fiber, since ***N***_tt_ has an eigenvalue in the parallel direction which is twice as large as the perpendicular directions). The mean square displacement of the fiber center is then 〈*r^2^*(*t*)〉 = 6*D_t_t*. Substituting the parameters in Table 1, we obtain 6*D_t_* ≈ 0.13 *μ*m^2^/s, and thus the time to diffuse a mesh size is given by 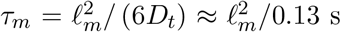. Since diffusion promotes mixing of the suspension and gives more opportunities for cross linking, our expectation is that thermal fluctuations should speed up the transition from the homogeneous meshwork to the composite bundle state, where bundles are made of a few fibers which must be close enough together to cross link. This assumes that the CL concentration is large enough for links to bind as soon as fibers are close enough together; we will analyze this assumption in Section 3.3.

To understand how thermal diffusion affects the bundling process, in Fig. 3 we plot the statistics both with (orange) and without (blue) thermal fluctuations. We see that the entire process is faster with diffusion, as we might expect, but the degree of acceleration changes before and after *τ_c_*. Before *τ_c_*, the process with diffusion is significantly faster than without; for instance, it takes about 3 seconds for 50% of the fibers to be in bundles with diffusion, while without diffusion it takes 12 seconds (Fig. 3 inset), which is a difference of a factor of 4. Indeed, the critical bundling time *τ_c_* ≈ 4 seconds with diffusion, while we have already seen *τ_c_* ≈ 16 seconds without diffusion, so that the difference is again a factor of 4. For *t* > *τ_c_*, when bundles start to coalesce, the difference is only a factor of 2; an exponential fit to the decaying bundle density gives a constant of 20 seconds for simulations with diffusion and 40 seconds for simulations without diffusion.

**Figure 3:**
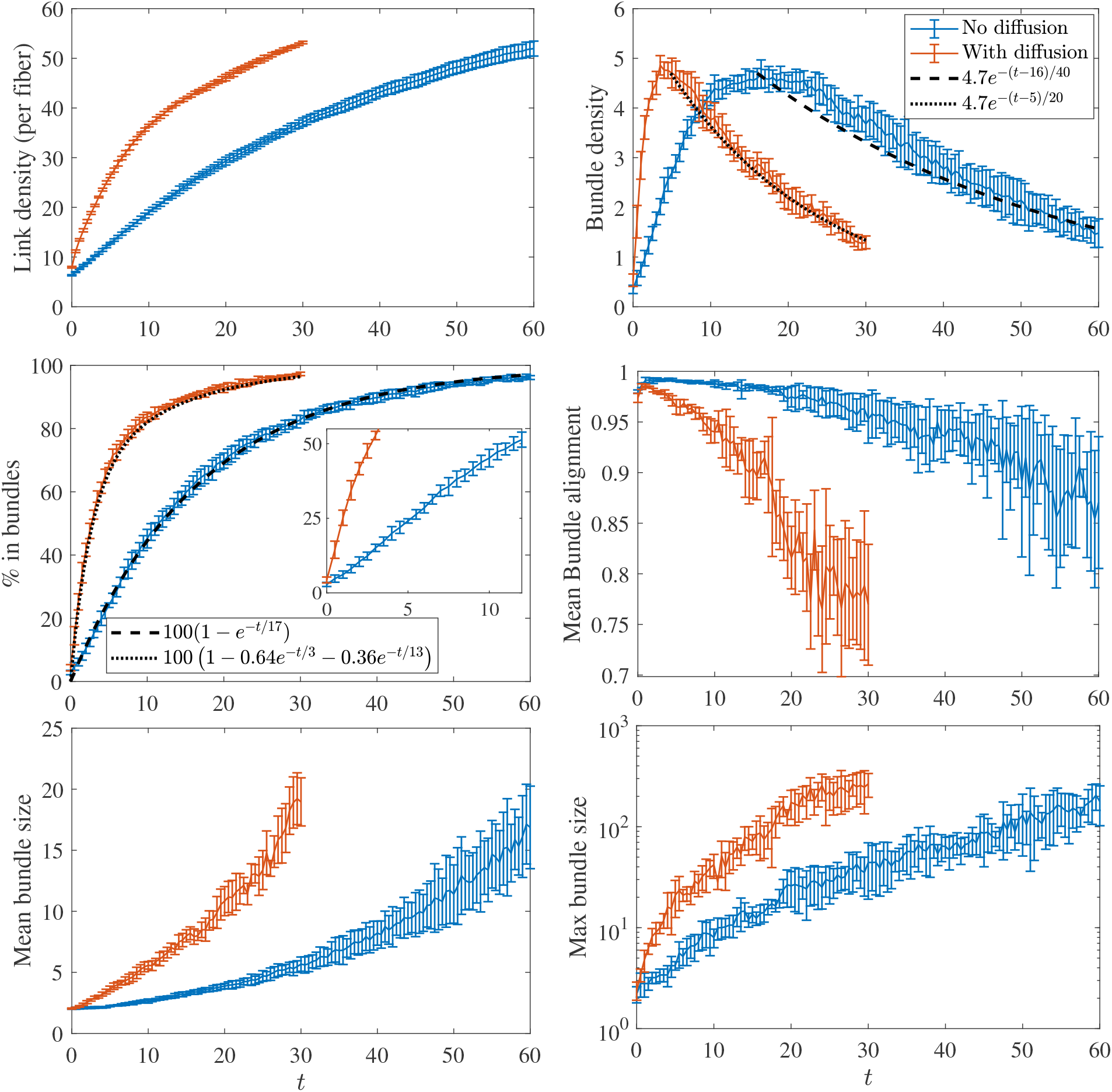
Statistics for the bundling process with and without thermal fluctuations. The blue lines show the results without thermal movement, while the orange lines show the results with translational and rotational diffusion. Here we use Δ*t* = 10^-4^; we have verified that the statistical noise exceeds the time stepping error for this time step size. The peak in the bundle density occurs at *τ_c_* ≈ 16 for systems without diffusion, while for systems with diffusion it occurs at *τ_c_* ≈ 4 seconds.

A similar relationship holds when we look not at the number of bundles (which depends on *F_B_*, the minimum number of fibers forming a bundle), but the percentage of fibers in bundles, which is independent of *F_B_* and shown in the middle left frame of Fig. 3. Unlike in the non-Brownian case, where a single timescale fits the data, the Brownian case requires two timescales for fitting, which are 3 seconds (which is close, but not equal to *τ_c_* ≈ 4 s), and 13 seconds. In this case, the new fast timescale of 3 seconds reflects the ability of *Brownian* filaments to freely diffuse translationally and rotationally early in the simulation. Later in the simulation, the filaments are arrested, and the timescale on which filaments enter bundles approaches that of non-Brownian filaments, 17 seconds. This provides more evidence for our two stage model of bundling, where thermal fluctuations make more of a difference in the first stage when fibers are less constrained by cross links. Sure enough, Fig. S4 (left, blue curve) shows that the mean squared displacement for simulations with Brownian motion decays exponentially to a constant, meaning that at times larger than *τ_c_* the Brownian motion is inhibited by cross linking, and therefore becomes less important. Equivalently, entropic effects (Brownian motion of fibers and cross-linker stretching) are more important at early times, while at later times energetic effects trap the fibers in the clustered bundle state.

To show that the network morphology has not changed when we add thermal movement, in Fig. S6 we show the networks at *t* ≈ *τ_c_* without and with thermal fluctuations. The composite bundle network morphology at *τ_c_* is similar between the two, which demonstrates that fluctuations speed up the pace of bundling without changing the types of bundles that evolve.

In subsequent sections, we will analyze how the timescale *τ_c_* that we use to quantify the speed of bundling depends on the microscopic parameters. While the precise value of *τ_c_* depends on the parameter *F_B_* (the minimum number of filaments in a “bundle”), Fig. 3 shows that this timescale can roughly capture the initial growth rate of the percentage of fibers in bundles, which is independent of *F_B_*. Since *τ_c_* is easier to measure by looking at the peak bundle density (and is in principle easier to measure experimentally through microscopy) than by fitting a double-exponential curve (which is an ill-conditioned problem for larger timescales), we will use the bundle density maximum as the definition of *τ_c_*. Of course, making *F_B_* larger increases *τ_c_*, as we show in Fig. S5 by setting *F_B_* = 5, but the ratio of *τ_c_* between the Brownian and non-Brownian system remains the same. However, increasing *F_B_* will cause us to miss the initial stage of bundling where two-filament bundles form and the fibers’ rotational degrees of freedom are arrested, so we will use *F_B_* = 2 henceforth.

### 3.3 Dependence of bundling timescale on actin and CL concentration

Our conclusion that thermal fluctuations significantly accelerate the initial stage of the bundling process is dependent on having a sufficient concentration of CLs. While thermal fluctuations undoubtedly increase the frequency of fibers coming close enough together for cross linking, the bundling process still must be initiated via binding of a CL. Consequently, in this section we consider a range of values of mesh size (actin concentration) and *k*_on_ (CL attachment rate, which is proportional to CL concentration), to get a more complete picture of how the critical bundling time *τ_c_* depends on these parameters. In particular, we will consider mesh sizes *ℓ_m_* = 0.2 (*F* = 675, *L_d_* = 3 *μ*m), 0.4 (*F* = 400, *L_d_* = 4 *μ*m), and 0.8 *μ*m (*F* = 338, *L_d_* = 6 *μ*m), and single-end binding rates *k*_on_ = 1.25, 5 (the base value), and 20 (*μ*m·s)^-1^. By changing the rate at which a single CL end binds to a fiber, we effectively vary the CL concentration.

Figure 4 shows the resulting evolution of the bundle density for the nine different systems, as well as the resulting critical bundling time *τ_c_*. For systems with large *k*_on_, where binding is essentially instantaneous once filaments come close enough together, *τ_c_* ≈ 3 s for the small enough mesh sizes of *ℓ_m_* = 0.2 and 0.4 *μ*m. Once the mesh size increases to 0.8 *μ*m, the bundling time increases, but only to about 4.5 s (see inset of Fig. 4). Thus *τ_c_* is not a strong function of mesh size for larger *k*_on_, which implies that the process for large *k*_on_, where there is always sufficient crosslinker available for binding, is primarily limited by cross-linking dynamics (alignment of filaments), with diffusion (across the mesh size) playing only a secondary role.

**Figure 4:**
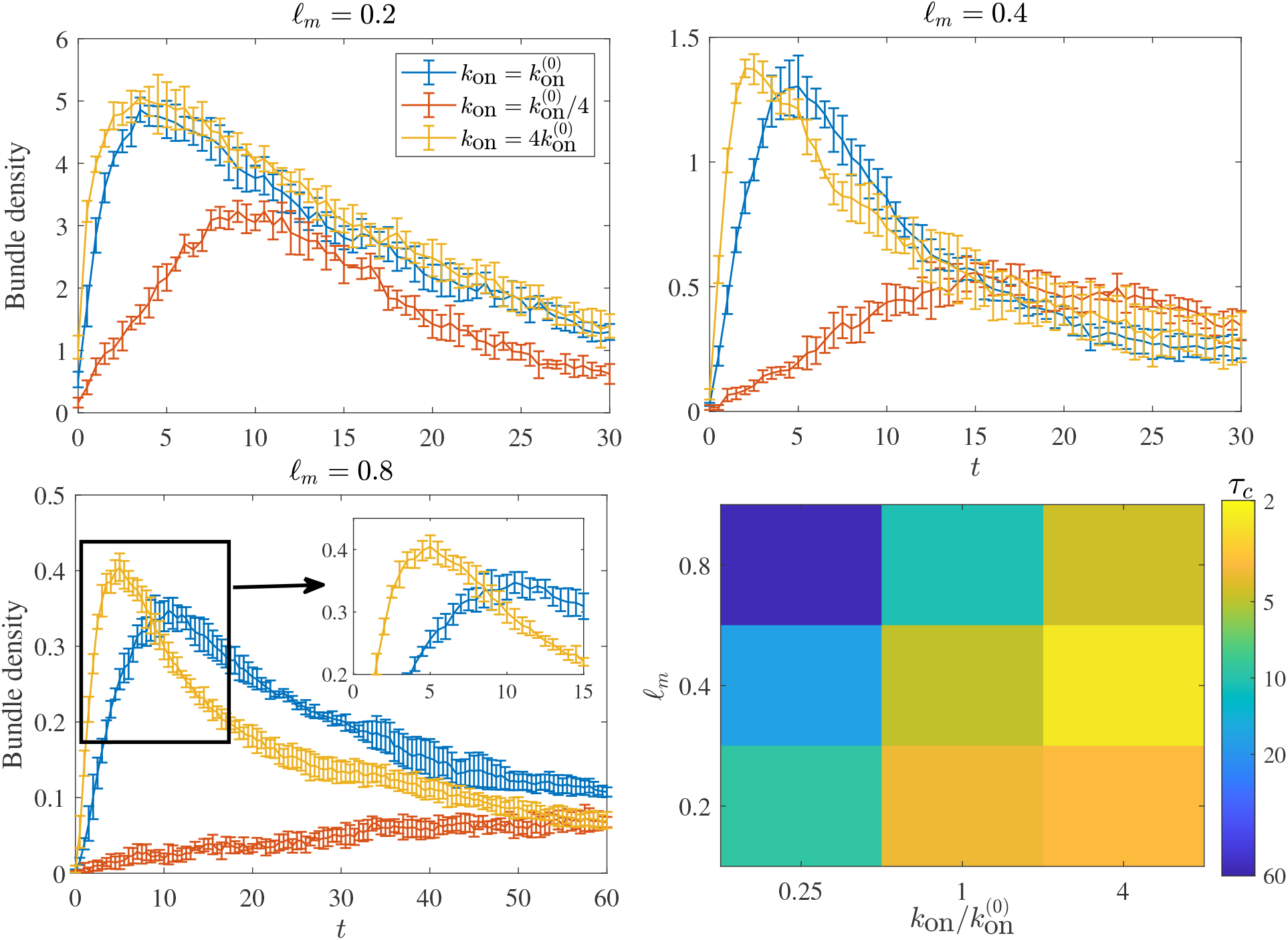
Bundling time scales for a range of initial mesh sizes *ℓ_m_* and binding rate *k*_on_. The first three frames show the trajectory of the bundle density for the different mesh sizes, where blue lines denote our base value of 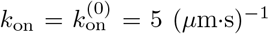, orange lines denote 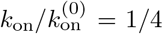, and yellow lines 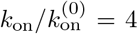. The bottom right frame shows the dependence of the critical bundling time *τ_c_* on *ℓ_m_* and *k*_on_.

Let us now consider the case of slower *k_on_*. In this case, filaments could come close enough to link together, but diffuse away before a CL can actually bind them. As a result of this, the bundling process is slowed, and in fact the peak bundle density drops. Indeed, as shown in Fig. 5, networks with smaller *k*_on_ (lower CL concentration) contain larger bundles at *t* = *τ_c_* than those with larger *k*_on_ (higher concentration). As shown in the supplementary videos, upon reducing *k*_on_, two filaments finding each other becomes the limiting step in the bundling process. This causes a slow growth of the bundle curve, and a bias towards larger bundles, which build up at a faster rate (relative to *τ_c_*), and the process is rate-limited by two-filament bundle formation. The scaling of *τ_c_* at small *k*_on_ (left column of the bottom right panel in Fig. 4) is reminiscent of a diffusion-limited process, as it increases from 9 s to 17 s to 56 s as the mesh size doubles, scaling approximately as 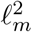 as the mesh size increases. In some sense, diffusion is actually a hindrance to bundling, since fibers that are close to each other diffuse away before a CL can bind them together.

**Figure 5:**
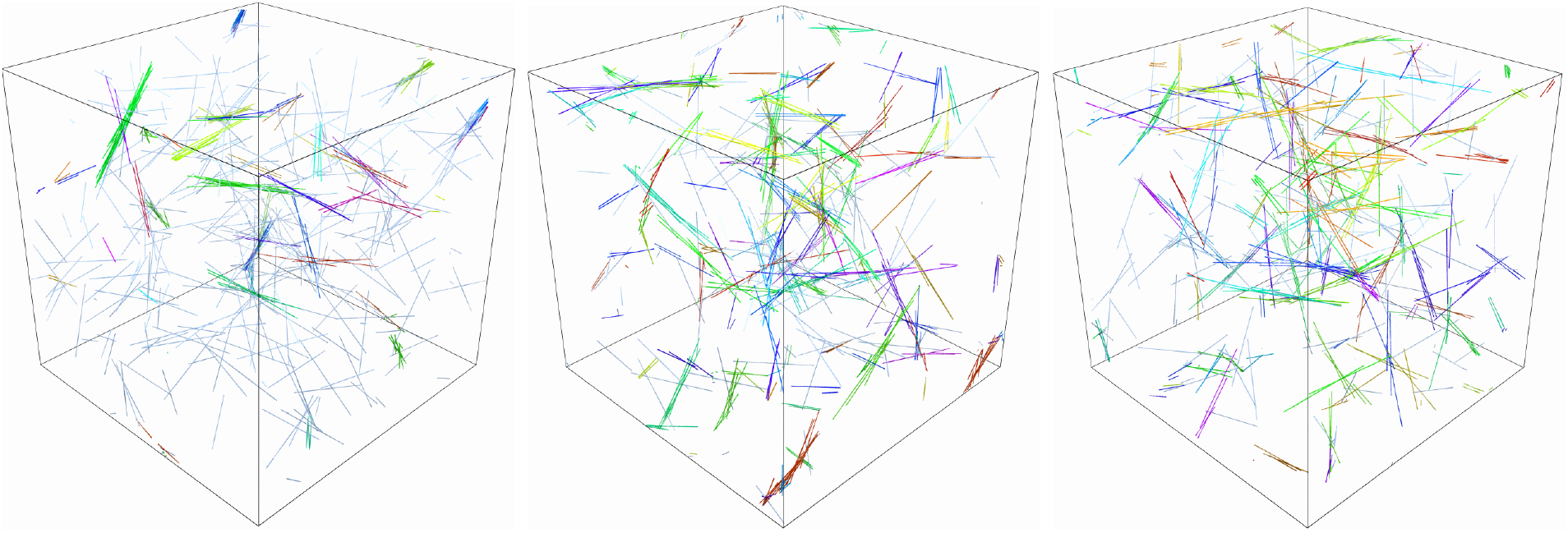
Snapshots of the network at *t* = *τ_c_* with initial mesh size *ℓ_m_* = 0.4 *μ*m (*F* = 400 filaments of length *L* = 1 in a domain of size *L_d_* = 4) with 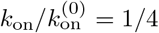 (left, *τ_c_* ≈ 17), 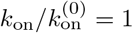 (middle, *τ_c_* ≈ 5), and 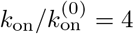 (right, *τ_c_* ≈ 2.5). A smaller *k*_on_ (smaller CL concentration) gives fewer but larger bundles at *t* = *τ_c_*, as well as a smaller percentage of fibers in bundles.

We note that a roughly constant bundling time can be achieved by decreasing *k*_on_ as the mesh size decreases (moving from the top right to the bottom left of the bottom right panel in Fig. 4). This implies that the relative concentration of CL required for a particular bundled state decreases with the mesh size, as has been found experimentally [40]. When the mesh size is smaller, the filaments are in contact for longer, and so it is less important that a CL be available immediately to bind them together. By contrast, filaments in larger-mesh-size systems are only in contact for a brief time, so relatively more CLs are necessary to ensure that these filaments are linked when they come into contact with each other.

### 3.4 Brownian motion is responsible for faster bundling with shorter filaments

We will now explore the dependence of the critical bundling time *τ_c_* on the fiber length. Experimentally, it has been shown that shorter filaments bundle faster [22, 11], but it is still unclear whether this is due to thermal movements, cross-linking kinetics, or some combination of both. In this section, we show that the experimental results can only be reproduced if we consider thermal movements, so that cross-linking kinetics are not responsible for the speedup in bundling. We use a fixed mesh size of *ℓ_m_* = 0.2 *μ*m, which translates to *F* = 675 filaments of length *L* = 1 *μ*m in a domain of size *L_d_* = 3 *μ*m, and *F* = 400 filaments of length *L* = 0.5 *μ*m in a domain of size *L_d_* = 2 *μ*m.

In Fig. 6, we show how the bundle density, percentage of fibers in bundles, and mean bundle size evolve for the two different filament lengths both (a) without and (b) with actin diffusion. In Fig. 6(a), we see that in the absence of Brownian motion the behavior in the two systems is similar, with the peak bundle density occurring in both cases around *τ_c_* ≈ 15 seconds. Furthermore, there is only a mild difference in the percentage of fibers in bundles over time. The mean bundle size is at most twice larger for the system with shorter filaments, but we would expect this since the filaments are twice as short and there are twice as many of them if *ℓ_m_* is fixed.

**Figure 6:**
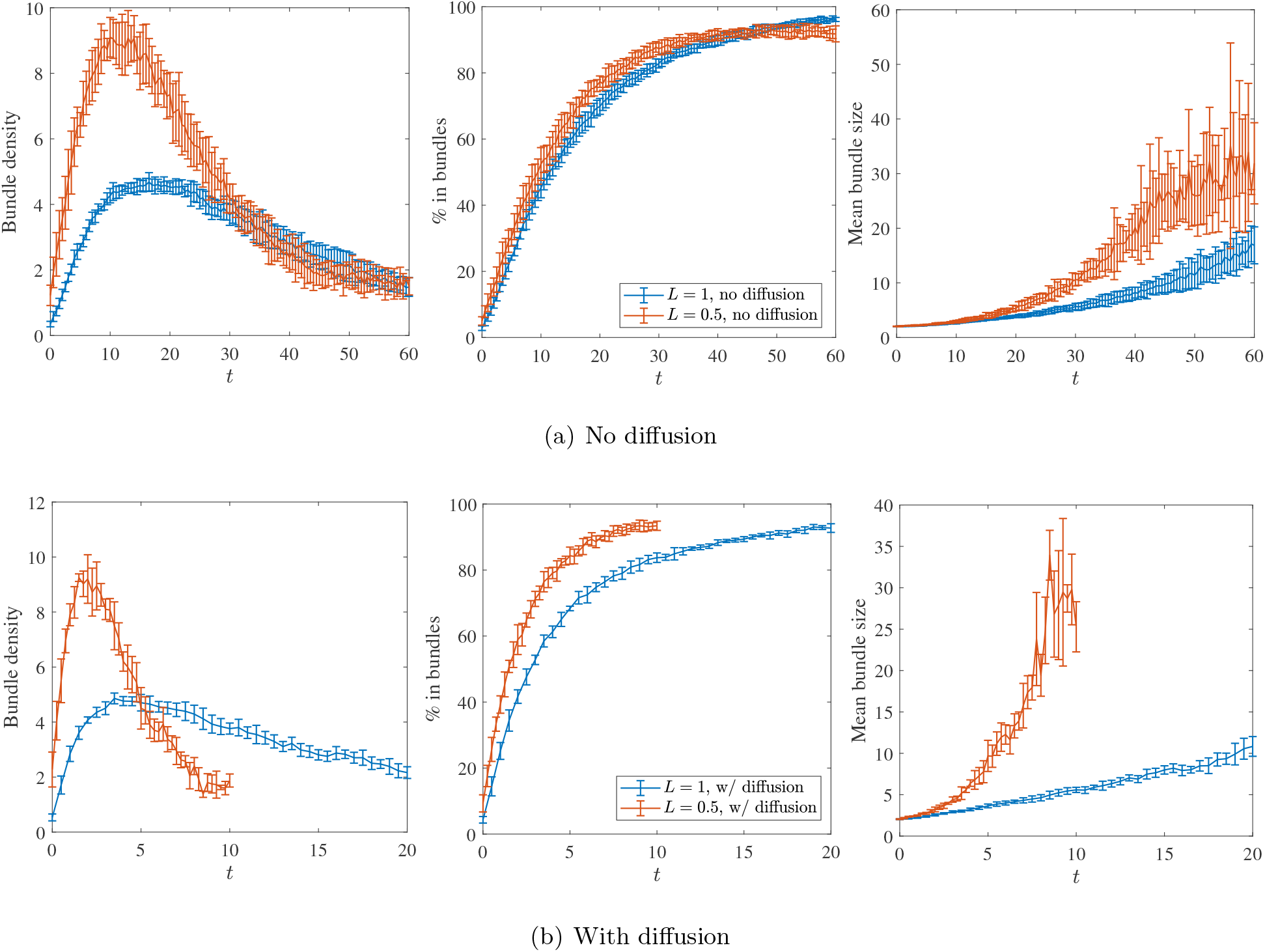
Effect of changing filament length for rigid fibers with and without Brownian motion, with constant initial mesh size *ℓ_m_* = 0.2 *μ*m. (a) Without fiber diffusion, we show the statistics for filaments of length *L* = 0.5 *μ*m (orange) and *L* = 1 *μ*m (blue), where we observe dynamics occurring on a similar timescale, especially in the initial stage (*t* ≤ *τ_c_* ≈ 20) of bundling. (b) When we add fiber diffusion, the bundling process for *L* = 0.5 *μ*m (orange) is significantly faster than *L* = 1 *μ*m (blue), because filaments can diffuse faster.

In Section 3.2, we showed that Brownian motion speeds up the bundling process by promoting mixing and more near-contacts of filaments. In particular, we saw that the time for a filament with length *L* = 1 *μ*m to diffuse a mesh size of *ℓ_m_* = 0.2 *μ*m is *τ_m_* ≈ 0.30 s, so that filaments can find each other rapidly and begin the bundling process. In the case of filaments with *L* = 0.5 *μ*m, our thermal diffusion coefficient (22) scales log-linearly with the fiber length, so that it takes *τ_m_* = 0.17 s to diffuse a mesh size of *ℓ_m_* = 0.2 *μ*m. We might expect, therefore, that at least the initial stages of the bundling process will be sped up by a factor of 2.

Figure 6(b) shows that this is indeed the case. For *ℓ_m_* = 0.2 *μ*m, the bundle density peak occurs around *τ_c_* ≈ 2 seconds when *L* = 0.5 *μ*m, while with *L* = 1 *μ*m it occurs around *τ_c_* ≈ 4 seconds, so it appears that bundling time with thermal motion scales linearly with filament length, which is in (approximate) accordance with the scaling of the translational diffusion coefficient. The faster bundling behavior also manifests itself in the link density and percentage of fibers in bundles, where we see that systems with shorter filaments reach a number of links or percentage of fibers about twice as fast. For instance, 80% of the fibers are in bundles by *t* ≈ 4 seconds in the *L* = 0.5 *μ*m case, while with *L* = 1 *μ*m the 80% mark is not reached until about *t* ≈ 8 seconds.

### 3.5 Ratio of bundling and turnover times control steady state morphology

Because we define the bundle density in terms of bundles of an arbitrary number of filaments (*F_B_* = 2), the precise value of the timescale *τ_c_* that we obtain is also somewhat arbitrary. Indeed, plotting the decay of the fibers’ MSD over the course of the simulation, as we do in Figs. S4 and S7, shows that *τ_c_* is not the only timescale characterizing the bundling process. However, if we increase the number of filaments required for a bundle to *F_B_* = 5, Fig. S5 shows that the peak in the bundle density occurs about a factor of 2 later in both Brownian and non-Brownian filament simulations. We therefore postulate that the *ratio* 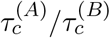 between systems A and B is a meaningful quantity, approximately independent of the definition of *τ_c_*, and can be used to predict the steady state morphology in systems with fiber turnover.

To test this, we introduce filament turnover with mean filament lifetime *τ_f_* (see [34] for implementation details) and fix *τ_f_* as a function of *τ_c_*, so that the ratio of the turnover times equals the ratio of the bundling times between the Brownian (B) and non-Brownian (NB) cases, 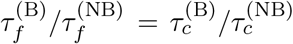, or, equivalently, 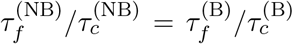. In Fig. 7 we vary the ratio *τ_f_*/*τ_c_* between 0.5 and 2 and plot the bundle density and percentage of fibers in bundles as they evolve to a steady state in each case. Despite the system of Brownian filaments having much faster bundling dynamics than the system of non-Brownian filaments, the morphology of the steady state is the same in the Brownian and non-Brownian cases, as is shown in the snapshots of Fig. S8.

**Figure 7:**
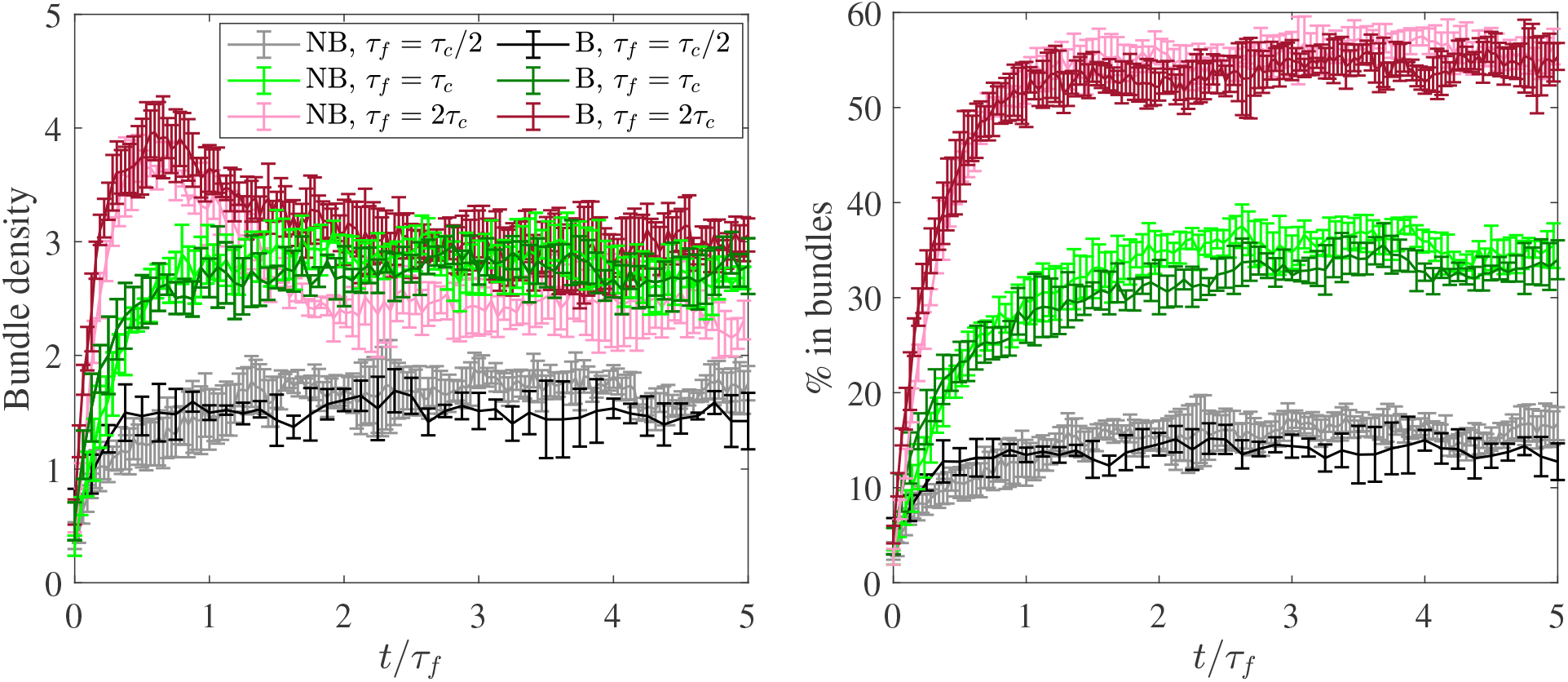
Steady state morphologies for systems with turnover. We introduce filament turnover with mean filament lifetime *τ_f_* (see [34] for implementation details) and observe the steady state bundle density (left) and percentage of fibers in bundles (right) for *τ_f_*/*τ_c_* = 1/2 (black), 1 (green), and 2 (red). Note that using a constant *τ_f_*/*τ_c_* in the two systems ensures 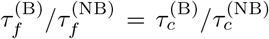. Using both non-Brownian (lighter colors, *τ_c_* ≈ 16 s) and Brownian (darker colors, *τ_c_* ≈ 4 s) filaments, we show that the steady state bundling statistics are roughly the same when *τ_f_*/*τ_c_* is matched.

## 4 Discussion

We used numerical simulations to investigate the kinetics of bundling in cross-linked actin suspensions. After validating that semiflexible actin fibers can be approximated as rigid in non-Brownian suspensions, we treated actin fibers as Brownian rigid, straight, slender rods, in accordance with a number of other simulation studies [12, 31]. We coarse grained the diffusion and binding and unbinding of *α*-actinin cross-linkers (CLs) into four microscopic rates: *k*_on_, *k*_on,s_, *k*_off_, and *k*_off,s_. This enabled the simulation of a gel with about 700 actin fibers and as many as 50 CLs bound to each fiber.

We found that, even without thermal movements, actin filaments can still bundle, as filaments that are initially close enough are linked together at small patches with CLs. These CLs pull fibers together and align them, thereby allowing more CLs to bind to other sections of the fibers. What results initially, for times smaller than the critical bundling coalescence time *τ_c_*, is a collection of bundles with a few highly aligned filaments, also called a composite bundle state [28]. For times larger than *τ_c_*, these bundles coalesce into larger bundles using a similar mechanism as that for individual fibers, and a clustered bundle state forms. Our critical bundling timescale *τ_c_* thus describes the initial time at which networks transition from the composite bundle state to the clustered bundle state. In networks with fiber turnover, a clustered bundle steady state is only possible if the turnover time is much larger than *τ_c_* [34]. Although our work leaves unclear the role of steric interactions in slowing down bundling, we did show that the strong cross-linking present at later times is sufficient to arrest the bundling process. In fact, strong cross-linking provides a force somewhat equivalent to steric interactions, since the finite rest length of the cross linkers keeps linked filaments apart [38, Note 23].

We quantified the role of diffusion throughout the bundling process, finding that it has a larger impact in the initial stages of bundling, when the filaments are not severely constrained by CLs and can move freely to find each other. We associated this stage with *t* < *τ_c_*, and showed that adding thermal fluctuations decreases *τ_c_* from 16 seconds to 4 seconds. We showed that the stage when bundles coalesce (*t* > *τ_c_*) is less affected by thermal diffusion (sped up by a factor of 2), since at that stage the filaments are constrained by CLs, which are more involved in bundle coalescence. This complements the observation in [11, 46] that bundling occurs faster in a fluid-like environment, where filaments can move freely prior to kinetic arrest.

At first glance, the order of magnitude of *τ_c_* that we obtained seems shorter than the characteristic bundling time obtained experimentally, which is generally reported to be on the order of minutes [36]. The comparison is difficult, however, since experimental times generally include polymerization, and the bundling timescale in experiments is defined by the onset of the clustered bundle steady state, which is much later than the composite bundle state where we define *τ_c_*. Nevertheless, the most instructive comparison is between our work and [11, Fig. 4], which shows experimentally that the addition of 10% nucleates (which speeds up the polymerization process) gives a saturated bundled state after 100 seconds of polymerization and bundling, where the bundles are made of at least 15 – 30 filaments and are spaced some 10 – 20 *μ*m apart. Given this observation, and the fact that bundling slows down over time, it is not difficult to imagine that the transition from the homogeneous state to the composite bundle state could take place on the order of 5 – 10 seconds after cross linkers are added to a system of (polymerized and capped) actin filaments.

While diffusion of fibers speeds up the bundling process, we showed that it must be combined with a sufficient concentration of CLs for rapid bundling to occur. In particular, we showed that a high concentration of CLs (high CL binding rate) can induce bundling for filaments of any mesh size, with a critical bundling time *τ_c_* that depends only weakly on the mesh size. By contrast, when the concentration of CLs is small, bundling is more difficult for any fixed mesh size, and gets near impossible as the mesh size increases, as near-fiber contacts become less frequent. This is in accordance with a number of experimental papers [40, 22] which find that bundling requires a critical CL concentration. In addition, because the fibers are in contact for a short time at larger mesh sizes, the system must be saturated with CLs for bundling to proceed at a reasonable rate. This saturation is less important at smaller mesh sizes, where fiber pairs come into contact more frequently. Translating our results to experimental parameters, we find that the ratio of the crosslinker concentration to the F-actin concentration that is needed for a particular bundling time scale decreases as the actin concentration increases, which is in accordance with existing experimental observations [40, Fig. 3].

As already mentioned, one of the drawbacks of some experimental studies is the sensitivity of the bundling time to the rate of actin polymerization. For example, it is shown in [11, Fig. 4(d)] that polymerization kinetics make an order of magnitude difference in the bundling kinetics. While simultaneous polymerization and bundling also occurs in vivo, our study here allowed us to divorce bundling and polymerization by focusing on a fixed filament length. By doing this, we showed that shorter filaments bundle faster exclusively because they can diffuse faster, because without thermal fluctuations we saw no difference in the bundling kinetics between short and long filaments. This clarifies why shorter actin filaments are able to associate more rapidly into bundles without the presence of a background actin mesh [22, 46].

There are, of course, other timescales that we could have examined in the bundling process. For instance, Figs. S4 and S7 show that the timescale for slow down of the fibers’ diffusivity, measured by the decay of their mean square displacement, is related to, but certainly not the same as, the critical bundling time *τ_c_*. Our choice to focus on the timescale *τ_c_* was motivated by our observation in previous work [34] that the steady state morphology of cross-linked actin networks is driven by a competition between bundling (which occurs on timescale *τ_c_*) and filament turnover (which occurs on timescale *τ_f_*). While it is intuitively obvious that increasing the turnover timescale *τ_f_* will produce a steady state with more bundles, it is fair to ask whether the ratio *τ_f_*/*τ_c_* alone controls the steady state morphology, or if some other microscopic parameters come into play. In Fig. 7, we showed that for turnover times *τ_f_* = *τ_c_*/2, *τ_c_*, and 2*τ_c_*, the gel evolves to a steady state where the bundle density and percentage of fibers in bundles depend primarily on the ratio *τ_f_*/*τ_c_*, for either Brownian or non-Brownian fibers (recall that *τ_c_* differs by a factor of 4 for these two cases). Snapshots in Fig. S8 show little qualitative difference between the network morphology of the Brownian and non-Brownian steady states for a fixed *τ_f_*/*τ_c_*. Thus, for a fixed turnover time *τ_f_*, the steady state morphology is controlled by *τ_c_*, which is the timescale we studied in detail here.

We can also extrapolate our results to the cell cytoskeleton, but this must be done with some caution because of the complexity of the in vivo system. The simulated actin network densities are characteristic of those observed in cell actin cortex, where mesh sizes are on the order 0.1 *μ*m [9]. Considering that the characteristic turnover times for the cell cortex are in the order of tens of seconds [13], longer than the characteristic bundling times our model predicts, the simple model prediction is that there is significant bundling in the cell cortex. However, to support this prediction, additional complexity, such as binding of filaments to the cell membrane and a mix of formin- and Arp2/3-generated filaments, will have to be added to the model. Similarly, in the future, the model could be modified to investigate effects of bundling rates that depend on mutual orientation of the filament pair [36].

Our study here used rigid filaments and coarse grained the dynamics of CL diffusion and binding. While we showed that *non-Brownian* semiflexible actin filaments can be approximated by rigid ones, we have not accounted for the transverse bending fluctuations in actin filaments. In some sense, softening the stiffness of the cross linkers, which gives a wider range of binding distances than might otherwise be possible, qualitatively accounts for this, but we plan to develop a numerical method that includes bending fluctuations in the future. We also hope to place our model of cross-linker dynamics on more rigorous footing by comparing it to a model that actually tracks the diffusion, binding, and unbinding of individual CLs. Other modeling studies addressed bundling in more complex systems, for example formation of unipolar bundles from a branched actin network [49] and bundling in the presence of a mix of CLs and myosin molecular motors [39, 3]. Interestingly, the appearance of the bundles in these more complex systems [39], which form when CL concentration is above a threshold value [5], resemble those predicted by our model without motors. Another level of complexity is limits on bundle sizes due chirality effects [17] and long-range electrostatic repulsion between the filaments (reviewed in [41]). Finally, in this work it was too difficult for us to simulate the experimental steady state clustered bundled morphologies, since we simulated actin filament lengths of 1 *μ*m and the observed steady states have bundles separated by hundreds of microns [40, 29]. More efficient, GPU-based, simulation techniques might enable the efficient simulation of even larger systems.

## Acknowledgements

This work was supported by the National Science Foundation through Research Training Group in Modeling and Simulation under award RTG/DMS-1646339 and through the Division of Mathematical Sciences award DMS-2052515. Ondrej Maxian is supported by the NSF via GRFP/DGE-1342536 and Alex Mogilner is supported by NSF grant DMS-1953430.

Code and input files for the simulations are available at https://github.com/stochasticHydroTools/ SlenderBody. Supplemental movies are available at https://cims.nyu.edu/~om759/BundlingVideos. All of our simulations were run on the NYU HPC Greene Supercomputer cluster.

## 5 Supplemental text

### 5.1 Rigid fibers as a special case of inextensible fibers

In [33], we discretized the inextensible system (2) by discretizing each fiber with *N* Chebyshev collocation points and representing the functions *α*_1_(*s*) and *α*_2_(*s*) by their Chebyshev coefficients, 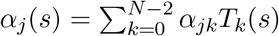. Our goal here is to show that setting *α_j_*(*s*) = *α_j_* = const. instead gives the straight rigid fiber kinematic operators (8) and (9). For convenience, we first restate the kinematic equations for inextensible fibers, which are Equations (41) and (44) in [33],

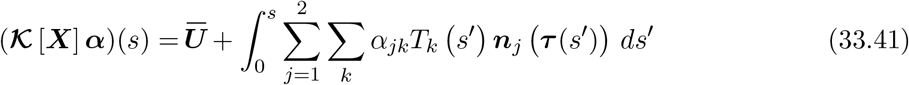

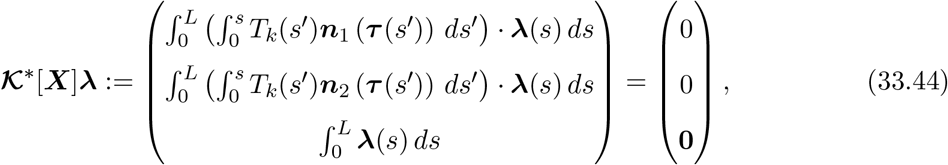

where the first two components of (33.44) hold for all *k*. Let us denote by 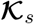 the operator 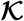 in the case of straight fibers with *k* = 0 being the only included Chebyshev mode, and likewise for 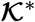. Then, since the fibers are straight, the orthonormal frame (***τ***, ***n**_i_*, ***n***_2_) is constant along the fiber, and thus 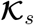 and 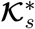 simplify to

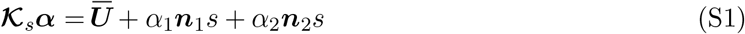

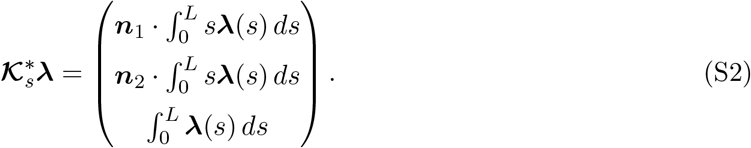

We now want to show that 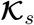 and 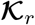 (defined in (8)) parameterize the same linear space of rigid motions. To do this, let us write ***X*** in (8) as an integral of the tangent vector

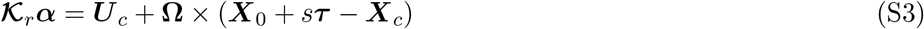

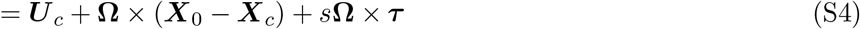

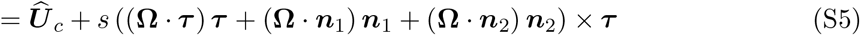

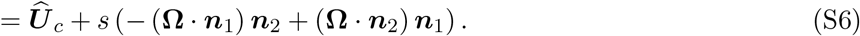

This is exactly the form of 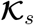 in (S1) with *α*_1_ = –**Ω** · ***n***_2_ and *α*_2_ = –**Ω** · ***n***_1_. So 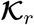 and 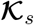 parameterize the same space.

To complete the equivalence, we now show that 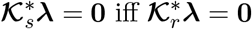. Obviously, 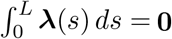 in both cases, so we only have to deal with the torque constraint. If we use the fact that 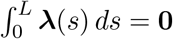, we can write the second component of (9) as

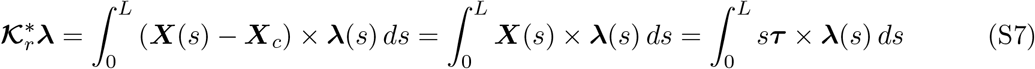

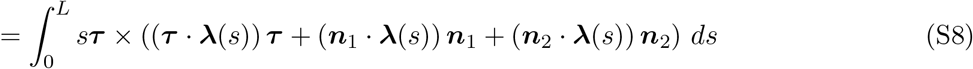

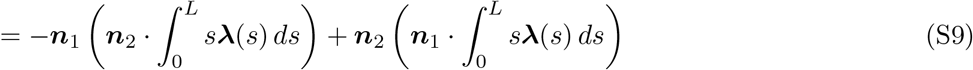

And now, since ***n***_1_ and ***n***_2_ are orthogonal and nonzero, we see from (S2) that 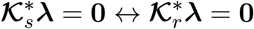. This shows that we can implement rigid fibers using the same algorithms as in [33], except we just need to keep a single (*k* = 0) Chebyshev polynomial. Note that the value of *κ* we use does not matter except for numerical stability since the fibers stay straight for all time, and so we set *κ* = 0.

### 5.2 Form and coefficients of the rigid body mobility matrix

Because of the symmetry of the fiber, the mobility matrix and its “square root” for a single fiber can be written in the form

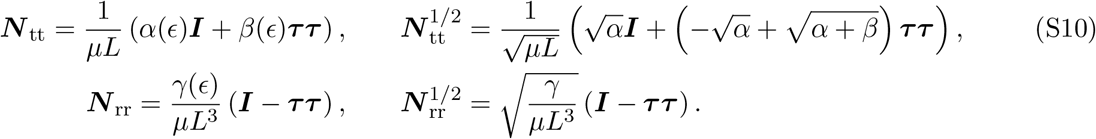

Notice that the rotational mobility has a null space of the tangent vector ***τ***. The dimensionless coefficients *α*, *β*, and *γ* are given for various *ε* in Table S1.

### 5.3 Temporal integrator for fluctuating fibers

In this section, we show that our temporal integrator for Brownian motion can accurately reproduce the steady state distribution of link strains. We place two parallel fibers a distance 0.05 apart, so that initially ***X***^(1)^(*s*) = (*s*, 0, 0) and ***X***^(2)^(*s*) = (*s*, 0.05, 0). At *t* = 0, the fibers are connected by a permanent CL attached at the point *s* = *L* = 1 on each fiber. We use rigid fibers with *N* = 50 points, CL variance *σ*/*L* = 0.005 (to simulate point-force-like springs), spring stiffness *K_c_* = 10 pN/*μ*m, and rest length *ℓ* = 0.05 *μ*m. We use *ε* = 0.004, *μ* = 0.1 Paos, and *L* = 1 *μ*m, as we do in most of the simulations in the main text. Because we are not interested in dynamics here, we use the local drag mobility, which is (4) without the integral term. The maximum stable time step is

**Table S1:** Mobility coefficients for the 6 ×6 rigid body mobility matrix ***N*** defined in (S10) for straight fibers. The numerical estimate of the matrix ***N*** is related to the slender body mobility ***M*** in (11), which is computed using intra-fiber hydrodynamics as discussed in Section 2.1.2.

| $\epsilon$ | $\alpha^{(\text{FP})}$ | $\beta^{(\text{FP})}$ | $\gamma^{(\text{FP})}$ |
| --- | --- | --- | --- |
| 0.01 | 0.3841 | 0.2230 | 3.4263 |
| 0.008 | 0.4020 | 0.2413 | 3.6412 |
| 0.006 | 0.4251 | 0.2646 | 3.9179 |
| 0.005 | 0.4396 | 0.2793 | 4.0931 |
| 0.004 | 0.4575 | 0.2973 | 4.3073 |
| 0.002 | 0.5129 | 0.3530 | 4.9721 |
| 0.001 | 0.5682 | 0.4085 | 5.6361 |

Δ*t* = 0.005 s, and so we will simulate both with Δ*t* = 0.0005 s (to get results with small temporal error) and Δ*t* = 0.0025 s (which is close to the stability limit). We simulate until *t* = 100 seconds in both cases and verify that we run for long enough that we have reached the steady state.

We expect the steady state probability density function (pdf) to be the Gibbs-Boltzmann distribution

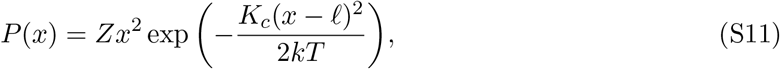

where the constant *Z* is chosen such that 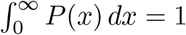, and the Jacobian *x*^2^ factor is necessary because *P*(*x*) is actually the one-dimensional analogue of the true three-dimensional distribution *P*(||*x*||). Figure S1 (left) shows that the steady state distribution with small Δ*t* agrees with the theory (S11). The right plot, which gives the distributions for Δ*t* =50% of the stability limit, shows that our temporal integrator can still reproduce the correct distribution with a larger time step size.

**Figure S1:**
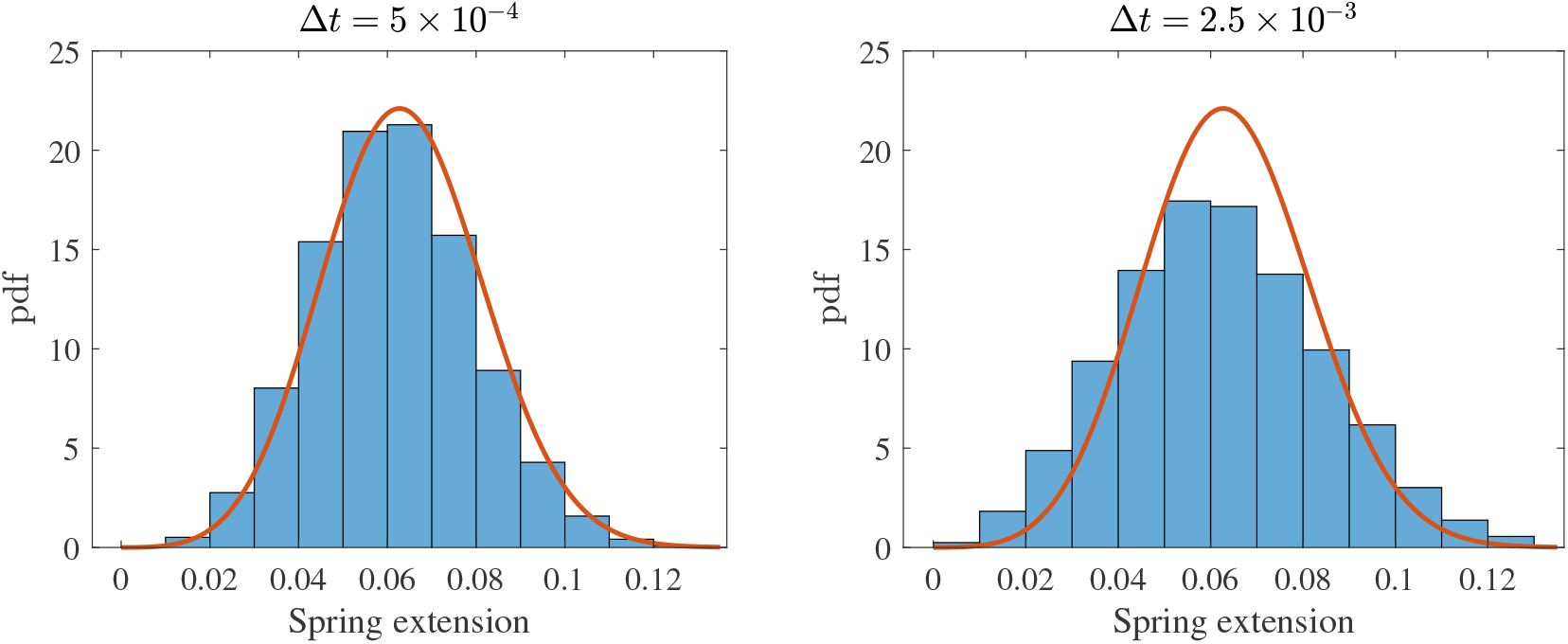
Steady state spring length distribution for two stiff fibers connected by a spring at their endpoint with rest length *ℓ* = 0.05 *μ*m. Histograms are the data, and red lines are (S11). Here we use rigid fibers with *N* = 50 points, CL variance *σ*/*L* = 0.005, and spring stiffness *K_c_* = 10 pN/*μ*m. We show Δ*t* = 5 × 10^-4^s =10% of the stability limit on the left, and Δ*t* = 2.5 × 10^-3^ s =50% of the stability limit on the right. The spring extension measurement is performed at the midpoint of the time step (after step 1 in Section 2.5).

**Figure S2:**
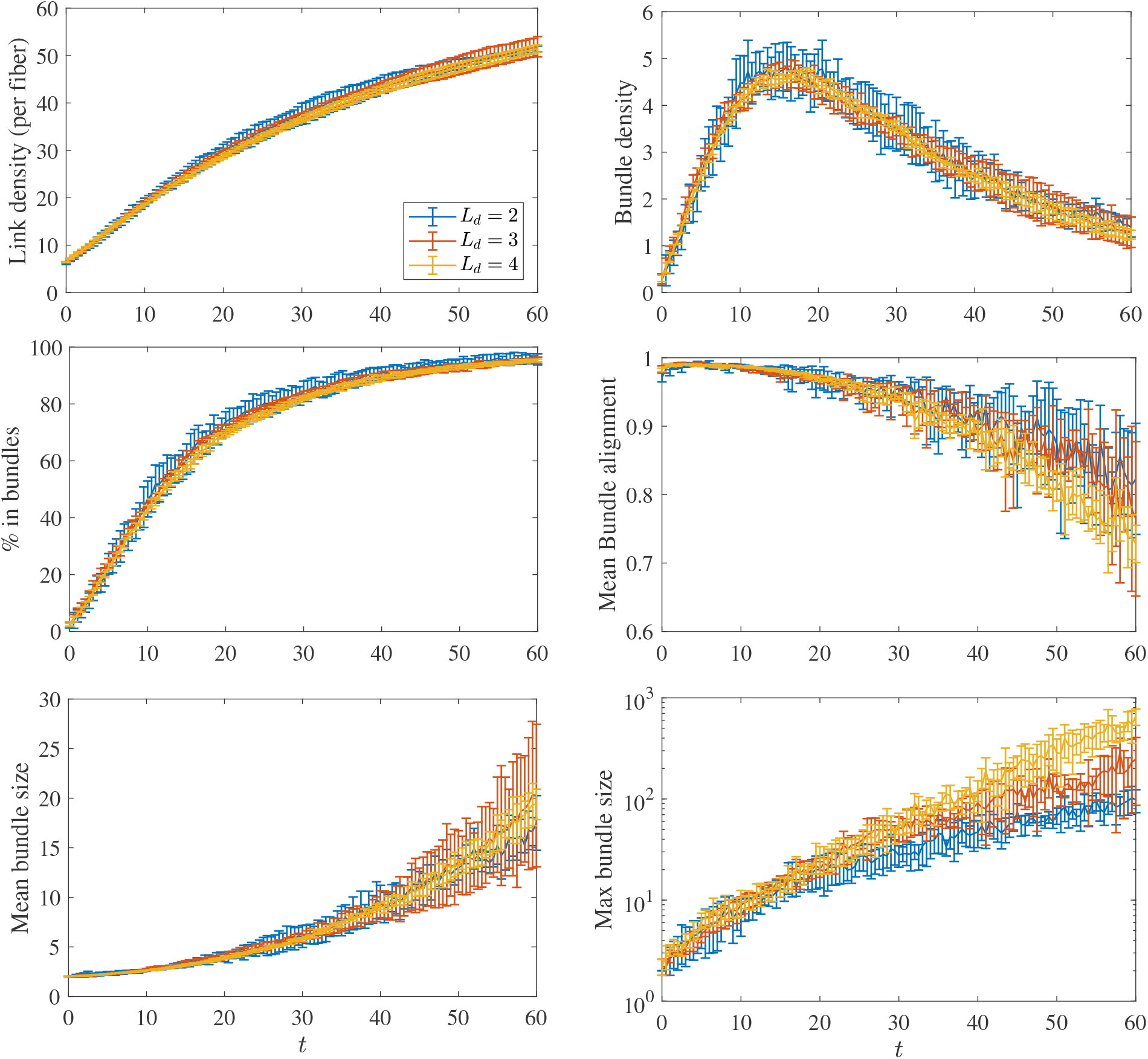
Link density, bundle density, % of fibers in bundles, mean bundle alignment, mean bundle size, and maximum bundle size, in a network of semiflexible (*κ* = 0.07 pN·*μ*m^2^) non-Brownian filaments with initial mesh size *ℓ_m_* = 0.2 *μ*m. We show curves with different domain sizes (in *μ*m) to establish that the statistics are repeatable in larger systems. The only statistic which is not repeatable is the maximum bundle size after many of the filaments have collapsed into one bundle (*t* ≳ 40 s).

**Figure S3:**
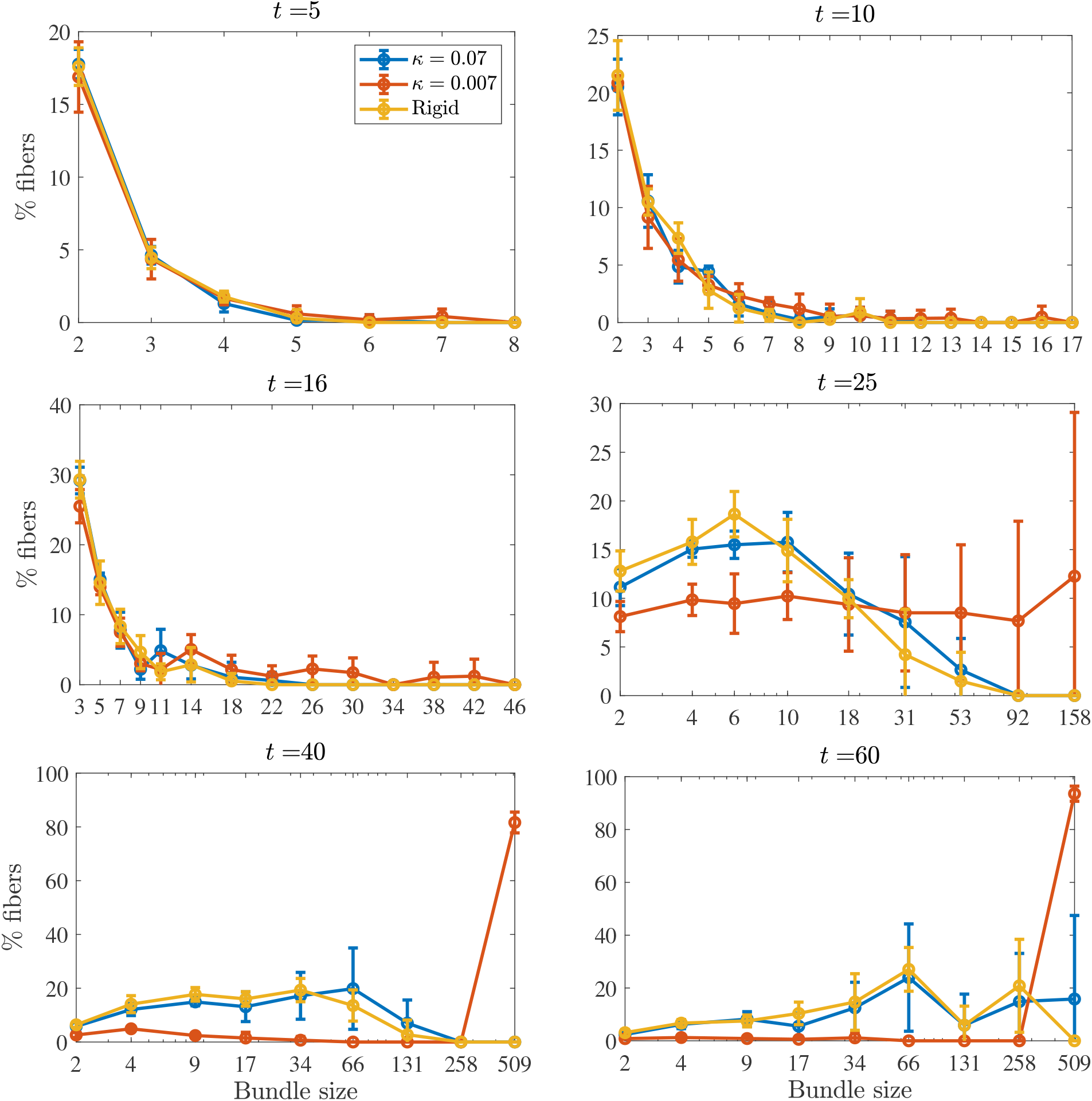
Bundle sizes over time in the system with *F* = 675 semiflexible filaments in a domain of size *L_d_* = 3 *μ*m (*τ_c_* ≈ 16). We show the percentage of fibers that are in bundles of various sizes over time for *κ* = 0.07 pN · *μ*m^2^ (blue), *κ* = 0.007 pN · *μ*m^2^ (orange), and rigid fibers (yellow). For times *t* = 25, 40, and 60 seconds, the *x* coordinate reflects the center of a histogram bin with logarithmically-scaled width. At *t* = 5 s, 25% of the filaments are in bundles of sizes 2 or 3, while most of the other fibers are not in bundles. At *t* = *τ_c_* = 16 s, about 50% of the fibers are in bundles of size 10 or less, with a small percentage in larger bundles, and the rest not in bundles at all (this is the composite bundle state). For semiflexible fibers with *κ* = 0.07 and rigid fibers, about 75% of the fibers are in bundles of size 30 or larger by *t* = 60, while for fibers with *κ* = 0.007 the entire suspension has coalesced together by *t* = 40 seconds.

**Figure S4:**
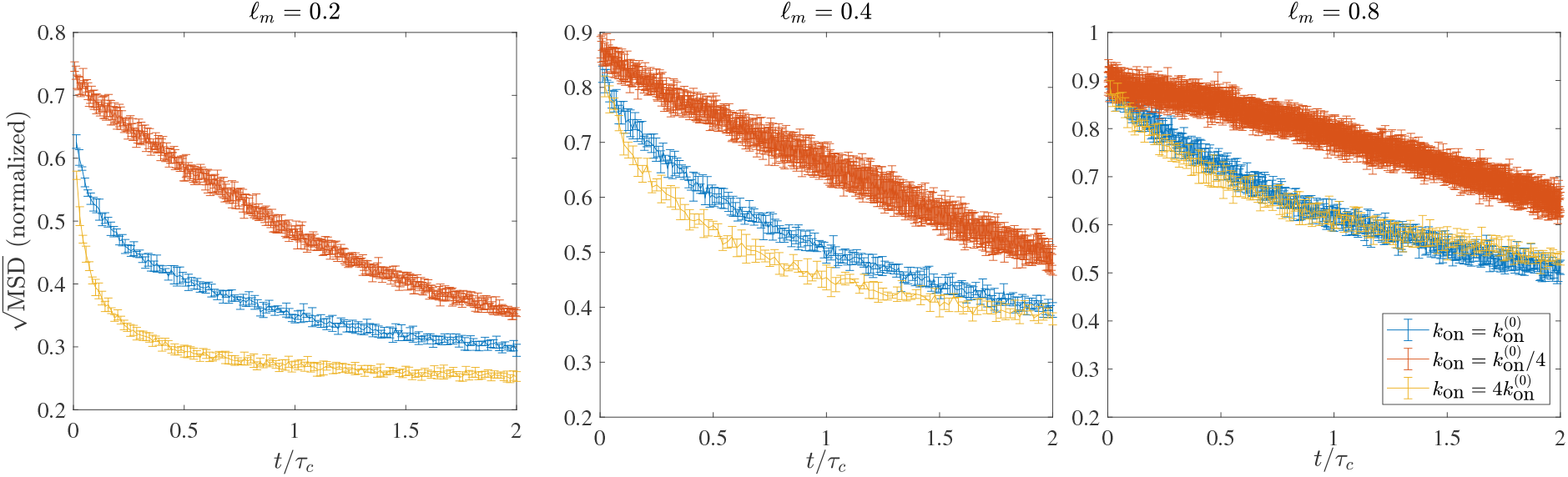
Square root of average mean square displacement of the fibers’ centers over a time span of 0.05 seconds for simulations with Brownian dynamics and varying *k*_on_ (colors) and mesh size (from small mesh size to large going from left to right), all normalized by the value for a freely diffusing fiber. We normalize time by *τ_c_*. While all of the curves show significant decay on the timescale *τ_c_*, it is clear that *τ_c_* is not the only timescale in the problem, since curves with small *k*_on_ are qualitatively different. This is not a surprise, since we saw in the main text that the bundling process with small kon is more biased towards large bundles (see Fig. 5).

**Figure S5:**
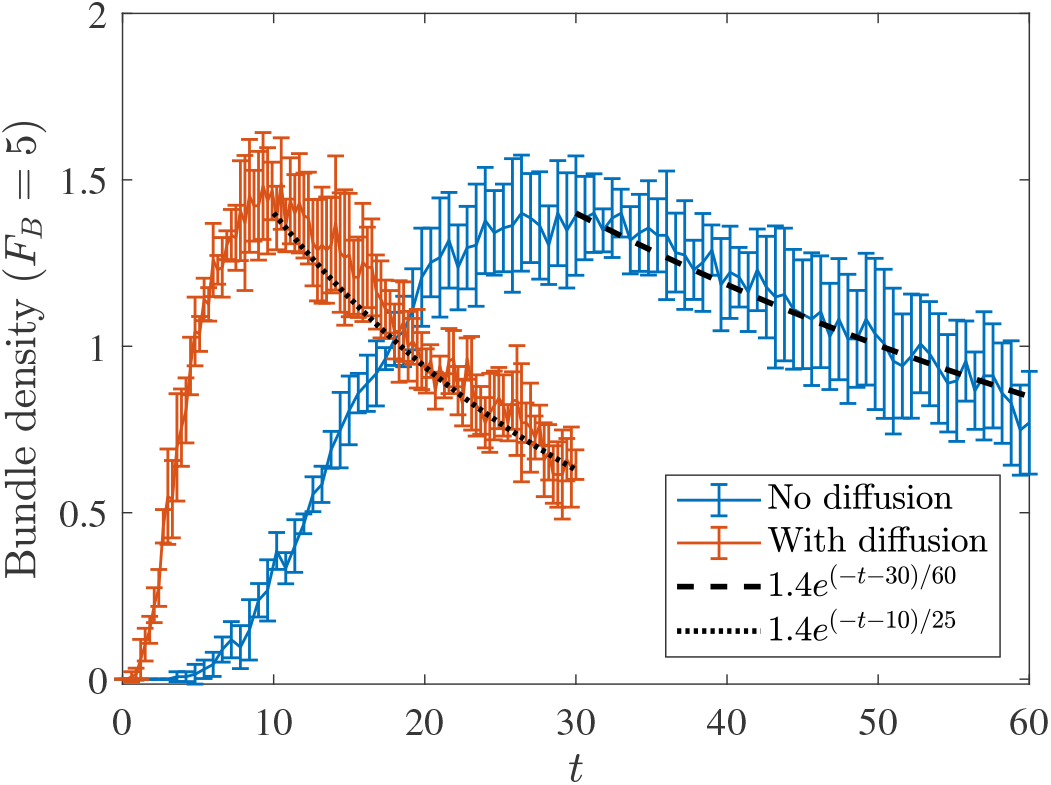
Comparing Brownian and non-Brownian filaments with a minimum of *F_B_* =5 filaments per bundle. In Fig. 3 in the main text, we used a definition of a bundle as having at least two filaments to conclude that Brownian motion accelerates the bundling process by more in the initial stages (factor of about 4) than in the latter (factor of about 2). When we increase to *F_B_* = 5 filaments per bundle, we observe the same characteristic growth and decay as with *F_B_* = 2, with the peak occurring three-fold faster in simulations with Brownian dynamics and the later dynamics being accelerated by a factor of about two (from a timescale of 60 s to 25 s). These confirm the qualitative (and quantitative) conclusions from Fig. 3 that we made in the main text.

**Figure S6:**
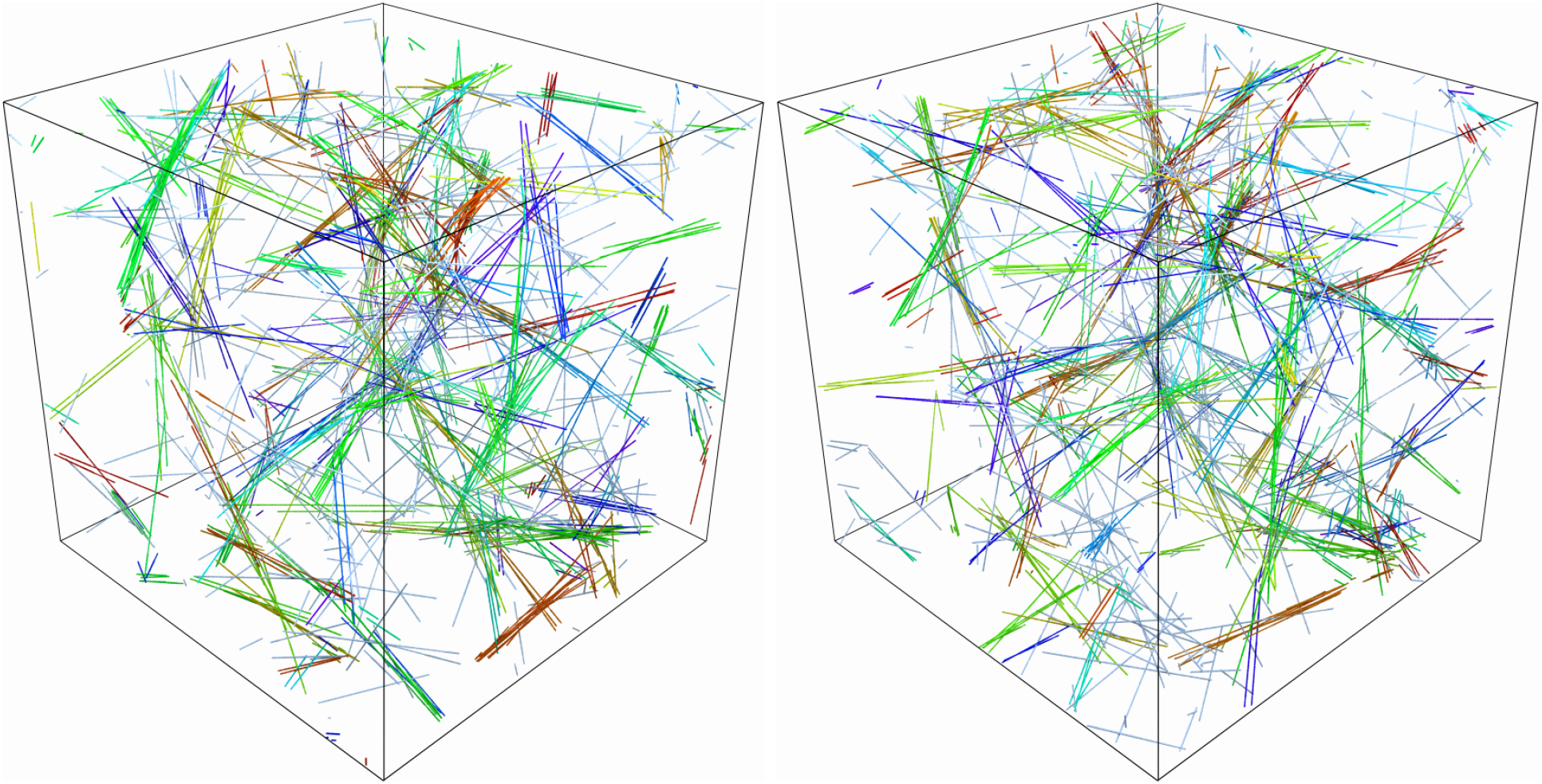
Representative snapshots of a network of rigid fibers without (left) and with (right) Brownian motion. Both snapshots are taken at *t* = *τ_c_*, which is 16 seconds for simulations without fluctuations and 4 seconds for simulations with fluctuations. The networks are qualitatively the same.

**Figure S7:**
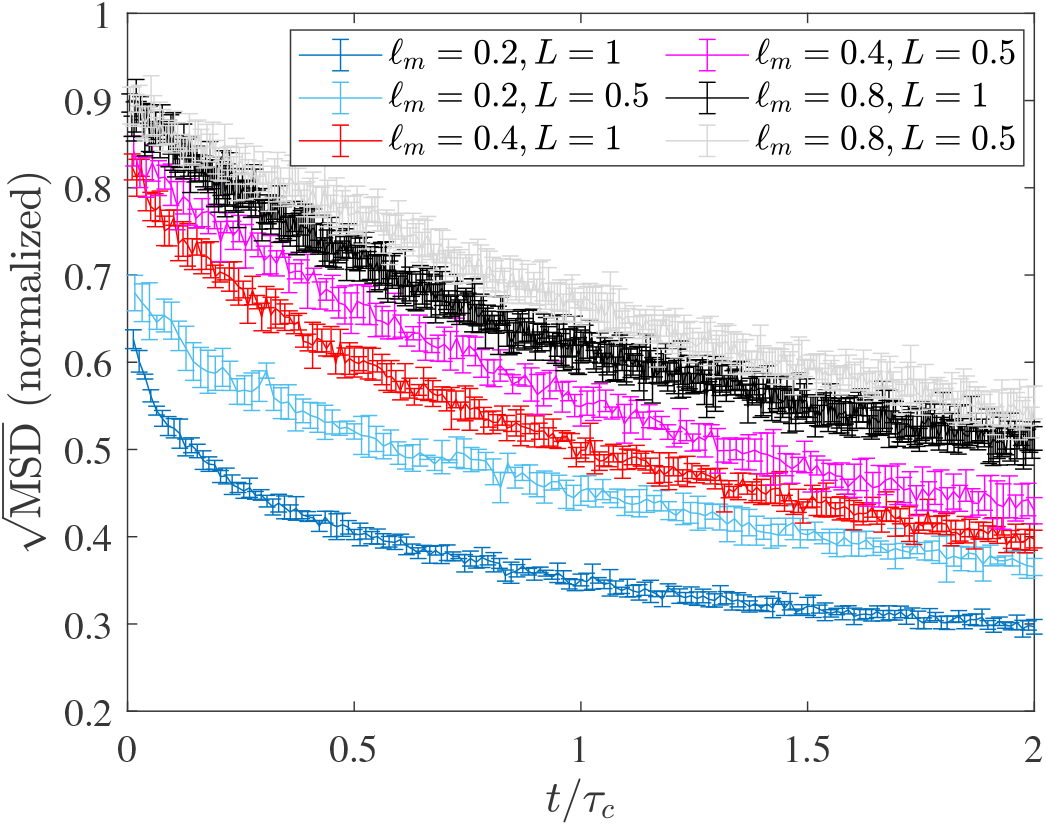
Decay of fibers’ centers’ displacement for different fiber lengths and mesh sizes. We show (the square root of) the average MSD of the fibers’ centers over a time span of 0.05 seconds, normalized by the free space diffusion for fibers of the same length. We show mesh size *ℓ_m_* = 0.2 *μ*m (blue), 0.4 *μ*m (red), and 0.8 *μ*m (black). Lighter colors are for filament length *L* = 0.5 *μ*m, darker are *L* =1 *μ*m.

**Figure S8:**
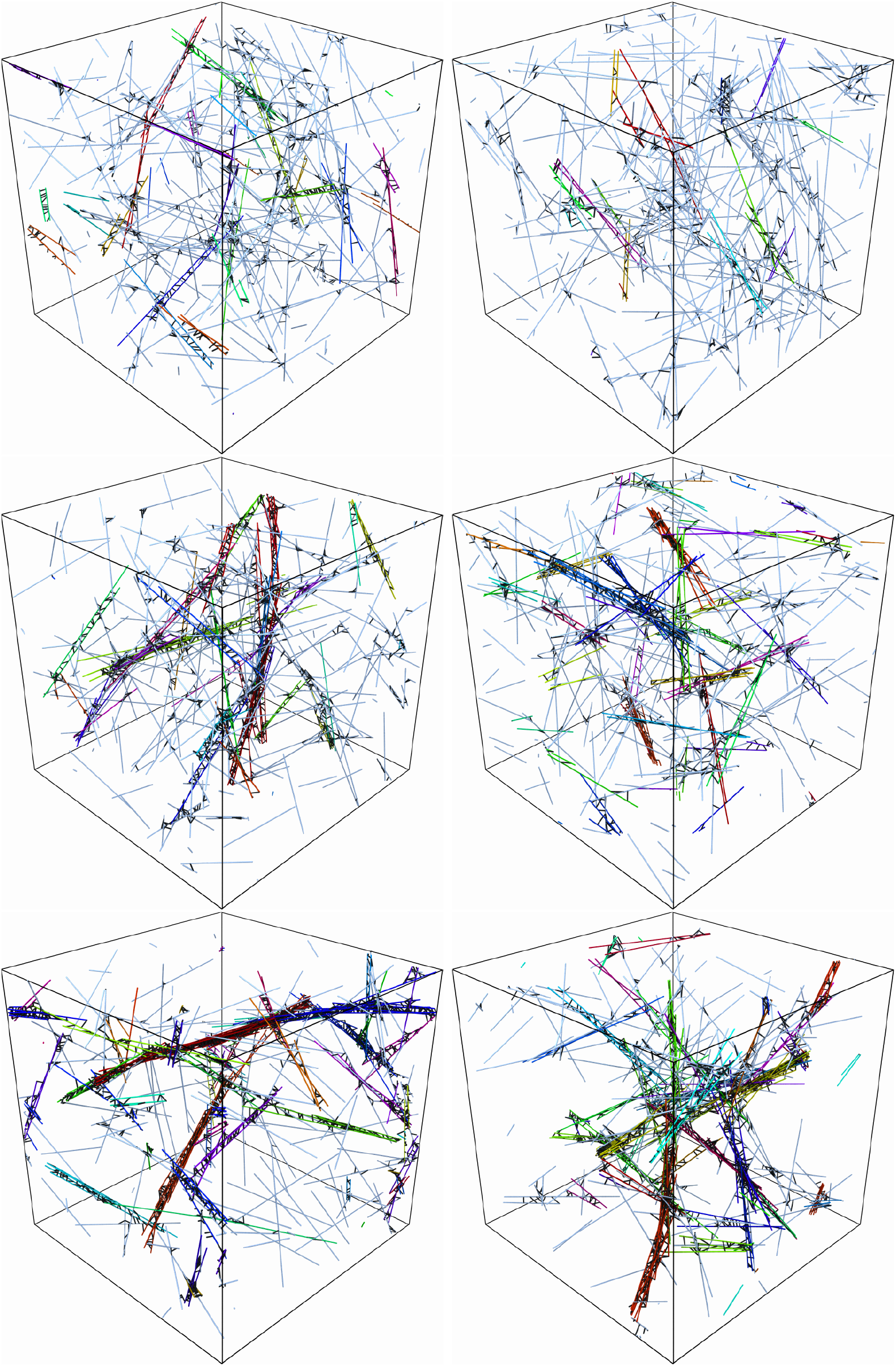
Steady states with fiber turnover for (top to bottom) *τ_f_* = *τ_c_*/2, *τ_f_* = *τ_c_*, and *τ_f_* = 2*τ_c_* for a system with *L_d_* = 2 and *ℓ_m_* = 0.2. The left column is for non-Brownian fibers (*τ_c_* = 16 seconds) and the right column is for Brownian ones (*τ_c_* = 4 seconds). There is little qualitative difference between the left and right columns, which indicates that the network morphology is controlled primarily by the ratio *τ_f_*/*τ_c_*.

## Footnotes

1 One can think of a bundle of *F_B_* filaments as contributing a weight ~ 1/*F_B_* to the bundle density, but a weight ~ *F_B_* to the percentage of fibers in bundles.

